# Human *APOE* allelic variants suppress the formation of diffuse and fibrillar Aβ deposits relative to mouse *Apoe* in transgenic mouse models of Alzheimer amyloidosis

**DOI:** 10.1101/2024.04.30.591932

**Authors:** Guilian Xu, Patricia Sacilotto, Carmelina Gorski, Parul Bali, Susan Fromholt, Quan Vo, Karen N McFarland, Qing Lu, David R Borchelt, Paramita Chakrabarty

**Affiliations:** Department of Neuroscience, University of Florida, Gainesville, FL-32610; Center for Translational Research in Neurodegenerative Disease, University of Florida, Gainesville, FL-32610; Department of Neurology, University of Florida, Gainesville, FL-32610; McKnight Brain Institute, University of Florida, Gainesville, FL-32610

**Author notes:** Address correspondence to Paramita Chakrabarty, 1275 Center Drive, BMS J484, Gainesville – 32610, USA., David R Borchelt, 1275 Center Drive, BMS J499, Gainesville – 32610, USA. Department of Pharmacology and Chemical Biology & Center for Neurodegenerative Disease, Emory University, Atlanta, GA-30322, USA. **Author emails:** Guilian Xu, Patricia Sacilotto, Carmelina Gorski, Parul Bali, Susan Fromholt, Quan Vo, Karen N McFarland, Qing Lu, David R Borchelt, Paramita Chakrabarty.

**Keywords:** Apolipoprotein E variants, seeding, aging, plaque morphology, amyloidogenicity

## Abstract

**Background:** Apolipoprotein E (apoE) modulates the deposition of amyloid β (Aβ) aggregates in Alzheimer’s disease (AD) in an isoform-dependent manner. In transgenic mouse models of AD-amyloidosis, replacing mouse *Apoe* alleles with human *APOE* variants suppresses fibrillar Aβ deposits. In the PD-APP transgenic mouse model, deletion of the *Apoe* gene led to selective reduction of fibrillar deposits with increased diffuse deposits. This finding suggested that apoE may have differential effects on different types of amyloid pathology.

**Methods:** Here, we investigated the interaction between the type of Aβ pathology in the brain and human apoE isoforms in different transgenic mouse models.

**Results:** In the APPsi model that develops predominantly diffuse Aβ plaques late in life, we determined that replacing mouse *Apoe* with human *APOE3* or *APOE4* genes potently suppressed diffuse amyloid formation, with apoE3 exhibiting a greater activity relative to apoE4. Relative to apoE4, apoE3 appeared to suppress Aβ deposition in the cerebral vasculature. In a second cohort, we accelerated the deposition of diffuse Aβ pathology by seeding, finding that seeded APPsi mice harboring *APOE4* or *APOE3* developed equal burdens of diffuse parenchymal Aβ. Finally, in the recently developed SAA-APP model that has a mix of dense-core and fibrous Aβ plaques, we found that replacing mouse apoE with human apoE suppressed deposition significantly, with the amyloid burden following the trend of *Apoe*>>*APOE4> APOE3*∼*APOE2*. In the SAA-APP and seeded APPsi models, we found evidence of apoE protein associated with Aβ plaques.

**Conclusions:** Overall, these observations demonstrate a capacity for human apoE to suppress the deposition of both diffuse and fibrillar-cored deposits, relative to mouse apoE. Notably, in the seeded paradigm, the suppressive activity of human apoE3 and apoE4 appeared to be overwhelmed. Taken together, this study demonstrates that *APOE* genotype influences the deposition of both cored-fibrillar and diffuse amyloid.

## Background

Apolipoprotein E (*APOE*) L4 is the major genetic risk factor of Alzheimer’s disease (AD), regulating the onset and progression of dementia in a gene dose-dependent manner (reviewed in [1,2,3]). Compared to the reference allele *APOE3*, *APOE4* is associated with higher plaque burden and earlier onset of disease (reviewed in [1,2,3]). The *APOE2* allele, on the other hand, extends lifespan and reduces the incidence of AD [1,2,3]. The mechanism is thought to be mediated via a combination of pathways that accelerate deposition of amyloid β (Aβ), induce Aβ oligomerization [4], disrupt Aβ clearance, modulate neuro-immune interactions [5,6] and influence Aβ-induced tau seeding [7].

Deposition of Aβ in the brain parenchyma and cerebral blood vessels is widely regarded as one of the initiating events for both familial and sporadic AD [8–12]. In AD patients, the proportion of dense core neuritic Aβ deposits correlates directly with immune abnormalities, neuronal network alterations and cognitive deficits [13]. On the other hand, diffuse Aβ plaques are less immune-reactive deposits that are also prevalent in cognitively normal aged humans [14]. Many proteins, including apoE, have been shown to co-deposit with Aβ [15], thus raising the possibility that these could have a role in the initial seeding phase or during subsequent plaque growth phase to facilitate the formation of neuritic plaques [16].

*APOE* genotype has emerged as a key modulator of Aβ deposition. Replacing mouse *Apoe* with human *APOE* in transgenic models of amyloidosis results in dramatic reductions in the burden of cored, thioflavin-positive, Aβ deposits [17,18]. Relative to mouse apoE, human apoE variants appear to be potent suppressors of cored Aβ deposition. Replacing mouse *Apoe* with human *APOE* variants in PD-APP mice resulted lower levels of cored Aβ deposits with the relative abundance stratifying as mouse *Apoe*>>*APOE4*>*APOE3*>APOE2 [19]. In the PD-APP mouse model of AD-amyloidosis, reductions in cored deposits were balanced by increases in the burden of diffuse deposits [20–22]. Paradoxically, eliminating both *Apoe* and *Apoj* in PD-APP mice led to substantial increases in both diffuse and cored Aβ deposits, leading the authors to conclude that in mice apoE and apoJ act cooperatively to suppress Aβ deposition [17]. In all three cases, diffuse amyloid deposits appeared to be less affected, suggesting that apoE may have a more significant impact on cored Aβ deposits rather than diffuse amyloid.

In the present study, we sought to specifically address the role of apoE in modulating the deposition of diffuse amyloid. We used a previously described *APP* transgenic mouse model (designated APPsi) that generates predominantly diffuse deposits [23,24]. When APPsi mice were crossed to *APOE3tr* and *APOE4tr* mice and aged to 21 months, we found that the presence of human apoE significantly suppressed the formation of diffuse amyloid deposits with APPsi mice expressing human apoE3 (APPsi/E3) having almost no deposition. The presence of human apoE3 was also a potent suppressor of Aβ deposition in cerebral vasculature. In a second cohort of APPsi/APOEtr bigenic mice where we induced pathology by intracerebral seeding [24,25], the presence of human apoE had little or no impact on seeding efficacy with both *APOE3* and *APOE4* genotypes showing similar burdens of diffuse amyloid. To compare these findings to mice with cored-fibrillar deposits, we crossed the APOEtr mice with the recently described SAA-APP knock-in model, that primarily develop fibrillar Aβ deposits [26], finding that the presence of human *APOE* alleles significantly suppressed the formation of cored-fibrillar deposits consistent with previous observations [19,21,22,27]. Overall, our study adds insights into the differential role of apoE isoforms in determining the severity and morphology of Aβ deposits in both progressively maturing and seeding-accelerated amyloidosis models.

## Materials and methods

### Mice

The PrP.APPsi mice express mouse APP-695 cDNA with a humanized Aβ sequence containing the FAD-associated APP-Swedish (KM670/671) and APP-Indiana (V717F) mutations from the mouse prion promoter [23]. The *SAA-APP* KI mice (B6.Cg-Apptm1.1Dnli/J; Jax Labs Stock # 034711) carries a humanized Aβ sequence with R684H, F681Y, and G676R mutations, along with the following human disease-associated mutations: KM670/671NL (Swedish), E693G (Arctic) and T714I (Austrian) [26]. The *WT-APP* KI mice, commonly known as B6J hAbeta (B6.Cg-Appem1Adiuj/J; Jax Labs Stock # 033013), carries the humanized Abeta1-42 region as in SAA-APP KI mice, without the disease-associated mutations. The APOEtr mice homozygous for human *APOE3* and *APOE4* alleles, via targeted replacement of the endogenous mouse *Apoe* gene, were obtained from Duke University under an MTA. All mice were housed in conventional air-filtered cages and maintained on standard diet and water. Mouse euthanasia was performed using approved protocols, following which one hemibrain was flash frozen and the corresponding hemibrain was drop-fixed in 4% paraformaldehyde overnight at 4°C. All procedures received prior approval from the Institutional Animal Care and Use Committee of the University of Florida and follow all applicable NIH guidelines.

### Preparation of brain homogenates and Intracerebral seeding

Frozen forebrain of a 13-mo PrP.APPsi mouse seeded with human AD patient homogenate (reported as AD#2 in [24]), which had been validated for robust induction of Aβ deposition, was homogenized in 10% (w/v) in sterile PBS by vortex and sonication (3 x 5 sec). The homogenate was then centrifuged at 3000 x g for 5 min at 4°C. Homogenates were immediately aliquoted and stored at -80°C until needed. 2µl of the 10% homogenate was delivered bilaterally into the cerebral ventricles of neonatal APPsi mice. Neonatal mice were cryoanesthetized and mouse brain lysates bilaterally using a 10 µL Hamilton syringe with a 33-gauge needle (Hamilton Company, Reno, NV). Mice were allowed to recover on a heating pad and transferred to their home cage. No adverse effects of injection were noted in previous study [24] and this study.

### Immunohistochemical, co-immunofluorescence and histology protocols

**Additional File 1:Table S1** provides information on different antibodies used in this study. 5-µm-thick sagittal sections of formalin-fixed paraffin-embedded (FFPE) hemibrains were used for immunohistochemical staining. Slides underwent de-paraffinization and rehydration in water, antigen retrieval in steam or citrate buffer as appropriate, 0.3% hydrogen peroxide treatment, and blocking in 3% serum, followed by primary antibody incubation at 4°C for 12-16 hr. Slides were then washed and incubated in secondary antibody (ImmPRESS Polymer Reagent, Vector Labs), and substrate development done using 3,3’-diaminobenzidine (DAB Peroxidase HRP Substrate Kit, Vector Labs). Slides were counterstained with hematoxylin, dehydrated and cover-slipped in mounting medium.

Co-immunofluorescence was done on deparaffinized slides that were blocked in 2% bovine serum albumin, incubated in primary antibodies overnight at 4°C and detected using Alexa fluor 488nm and Alexa-fluor 594nm conjugated secondary antibodies. Slides were washed and mounted in Fluoromount and imaged using Keyence BZ-X imager.

Campbell-Switzer Silver Stain was done according to a detailed protocol provided by Dr. Switzer (NeuroScience Associates, Knoxville, TN) [28].

Thioflavin S staining was done on FFPE hemibrain sections using 0.02% Thioflavin S solution according to Guntern modification of standard protocols [29]. Slides were deparaffinized and hydrated before background fluorescence was quenched using 0.25% Potassium permanganate, followed by 1% Potassium metabisulphite and 1% oxalic acid. Slides were then stained with Thioflavin S for 7 minutes, followed by quick differentiation in alcohol. Slides were washed and mounted in Fluoromount.

### Aβ analysis methods

To quantify Aβ deposit burden in APPsi, APPsi/APOEtr, SAA-APP or SAA-APP/APOEtr mice, we used FFPE slides cut at a level of approximately 1mm lateral to midline. Three sections from each mouse, spaced 20µm apart, were immuno-stained with Aβ antibody 33.1.1 and sections were scanned in an Aperio slide scanner. The percent immunoreactivity for each region was computed using the Aperio Positive Pixel Count program (Aperio, Vista, CA, USA). The burden of deposits for each slide in each cohort was analyzed in GraphPad (Prism 10, version 10.1.2), using 1-way ANOVA with corrections for repeated measures.

### Scoring of Cerebral CAA

An experienced observer unbiased towards outcome scored the CAA burden on silver-stained slides based on the following criteria: 3 = numerous stained parenchymal vessels per section; 2 = frequent stained vessels per section; 1 = limited number of stained vessels per section; 0 = no stained vessels per section.

### ELISA determination of Aβ

For the APPsi mice, frozen hemibrains were pulverized, weighed and extracted serially in RIPA buffer (50mM Tris-HCL, 150mM NaCl, 1% Triton 0, 0.5% Deoxycholate, 0.1% SDS), 2% SDS and 70% formic acid as described previously [30]. For the *SAA-APP* mice, frozen brains were first extracted in sterile PBS, followed by extraction in 70% formic acid. For capture of Aβ, 96 well Immulon 4 HBX plates (Thermoscientific) were coated with either 13.1.1 antibody (for Aβ40) or 2.1.3 antibody (for Aβ42) at a concentration of 20µg/µl. Brain homogenates were applied at different dilutions for interaction with Aβ antibodies at 4°C overnight. Detection was performed using HRP-conjugated 33.1.1 antibody. Aβ standards obtained from Millipore Sigma were prepared according to manufacturer’s instruction. The plates were visualized using SpectraMax device (Molecular Devices) and results were analyzed using SoftMax software (Molecular Devices).

### Immunoblotting for apoE and APP

RIPA homogenates were separated on 4-20% Tris-glycine gel (Novex, Invitrogen). Proteins were transferred to PVDF membranes, blocked for 1 hour in 0.5% casein and incubated overnight at 4°C in primary antibody. Detection of signal was performed using multiplex Li-Cor Odyssey Infrared Imaging System (Li-Cor Biosciences, Lincoln, NE, USA). Band intensities were quantified and standardized using ImageJ Software (NIH).

### Determination of AD transcriptome signature

RNA was extracted from frozen hemi forebrains using miniprep column (Invitrogen) purification of Trizol homogenates. 100□ng of the total RNA was analyzed on a commercially available NanoString Neurodegeneration codeset (NanoString, USA). Raw count data was checked for data quality using nSolver version 4.0 and imported into R version 3.6 and Bioconductor version 3.9. To calculate the individual gene expression signatures, geometric means of a set of genes characterizing each profile were used as described before [31].

### Statistics

All statistics have been done using 1-way Anova with Tukey’s test when appropriate or unpaired 2-tailed t-test as indicated in the Figure Legend.

## Results

### Human apoE isoforms suppress amyloid deposition in a model of diffuse Aβ pathology

Numerous studies have established that apoE is a potent modifier of AD-associated amyloid pathology (reviewed in [32,33]), specifically that apoE4 isoform worsens Aβ pathology relative to apoE3 isoform. Studies in transgenic mice have demonstrated that apoE expression at early phases of amyloid coalescence is critical in amyloidogenesis [34,35]. In studies in which mouse *Apoe* alleles have been replaced by human *APOE* alleles, investigators have noted that relative to mouse apoE, the presence of human apoE suppresses the formation of cored-fibrillar deposits [19,36]. We used three independent cohorts of Aβ deposition scenarios to ask the question of how the presence of human apoE or the mouse apoE isoforms drives Aβ pathology in progressively maturing and seeding-accelerated models.

In the first experiment, we examined the influence of apoE on the PrP.APPsi model (‘APPsi’) model, which develops primarily diffuse Aβ deposits beginning at 15-mo of age [23]. The APPsi mice were crossed to *APOE3tr* and *APOE4tr* mice to produce mice that were hetero- or homozygous for each *APOE* allele. Our purpose was to determine the influence of human apoE (APPsi/E3 and APPsi/E4) and mouse apoE (APPsi/e) on the natural evolution of Aβ deposition by aging these cohorts to 21 months of age. To control for the influence of strain background, we crossed APPsi mice to C57BL/6J mice for 4 to 5 generations to mimic the background of APPsi mice crossed to B6 congenic *APOE3tr* and *APOE4tr* mice. Using a neuropathological scoring criteria score, we observed that 21-mo APPsi/E3 mice and APPsi/E4 mice had almost no amyloid deposition in the cortex and hippocampus as compared to APPsi/e mice (**Fig. 1a-d**; p<0.0001 relative to APPsi/E3 and APPsi/E4 mice; **Additional File 1:Suppl. Table S2**). Mice that were heterozygous for human *APOE3* or human *APOE4* (carrying one copy of human *APOE* and one copy of mouse *Apoe*) displayed higher amyloid scores relative to mice homozygous for the corresponding human allele (**Additional File 1:Suppl. Fig. S1**; p<0.01 for both *APOE3* and *APOE4*; **Additional File 1:Suppl. Table S3**). This finding indicates that human apoE suppresses the deposition of diffuse amyloid. Consistent with our previous reports, aged APPsi/e mice developed prominent vascular amyloid staining (**Fig. S2a**) [23–25]. Neuropathological scoring of silver-stained sections revealed that both APPsi/e and APPsi/E4 mice had higher incidence of vascular Aβ deposition compared to APPsi/E3 mice (p<0.0001 for APPsi/e vs. APPsi/E3) (**Fig. 1e-f**; **Additional File 1:Supplementary Table S2**). Co-immunofluorescence analysis revealed only limited association of apoE with the diffuse Aβ deposits in these aged mice (**Additional File 1:Fig. S2b**).

**Figure 1.**
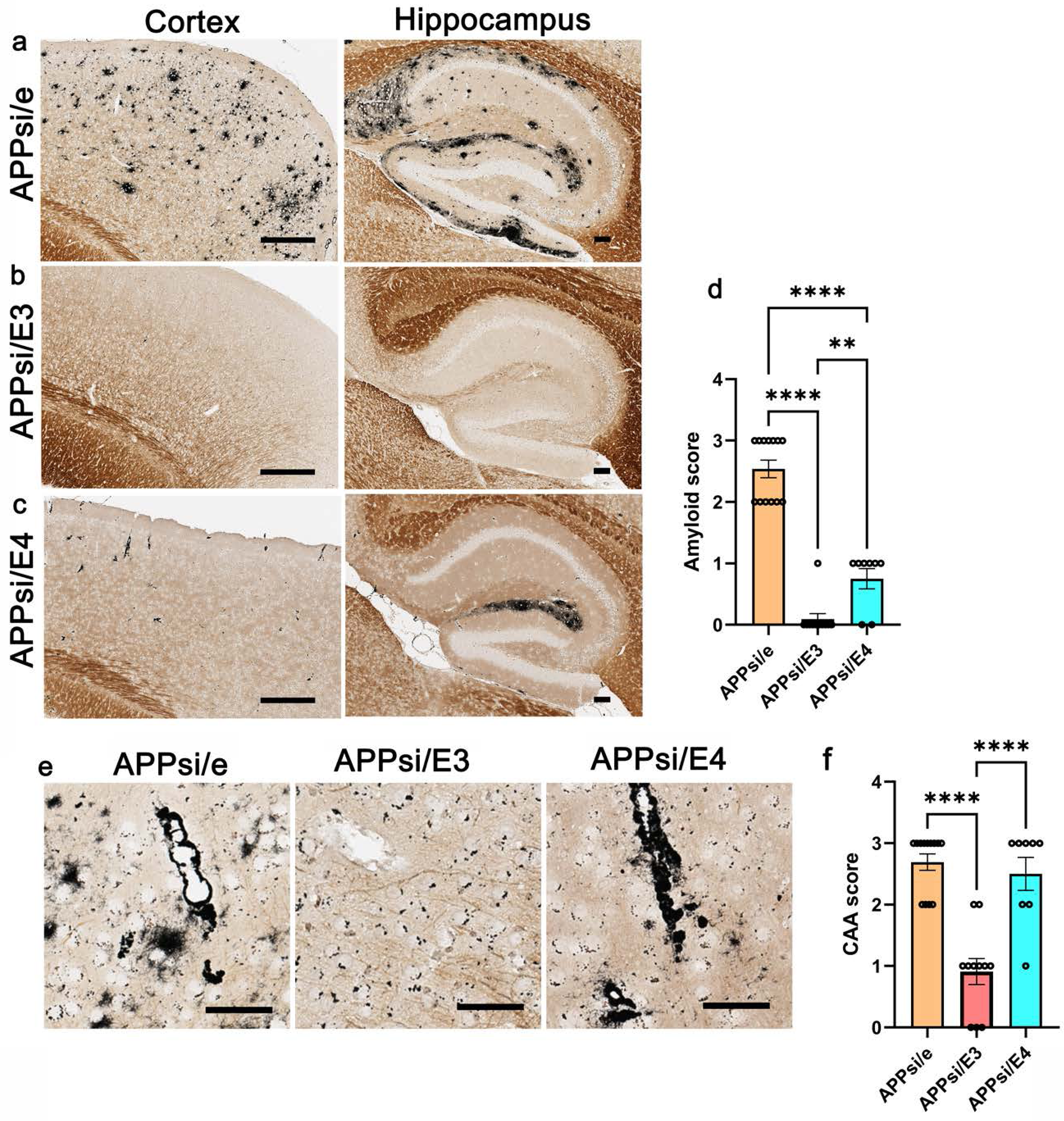
Characterization of aged APPsi mice expressing mouse and human apoE isoforms. **a-d.** FFPE brain sections of 21-mo old bigenic APPsi mice that are homozygous for mouse *Apoe*, or human *APOE3* and *APOE4* were stained with Campbell-Switzer silver stain to visualize Aβ plaques in the forebrain, hippocampus and cortex. Amyloid scoring was done by an experienced observer. Scale of 0 (none), 1 (low), 2 (medium) and 3 (highest). Scale bar, 100 µm. **e-f.** Representative silver-stained FFPE sections showing cortical CAA in these cohorts and burden count analysis. Scale of 0 (none), 1 (low), 2 (medium) and 3 (highest). Scale bar, 50 µm. n= 13 for APPsi/e, 11 for APPsi/E3 and n=8 for APPsi/E4 mice. 1-way Anova with Tukey’s test, ****p<0.0001.

We further performed immunohistochemical determination of Aβ burden using an N-terminal specific anti-Aβ antibody (**Fig. 2**). Analysis of immunostained Aβ burden revealed a pattern consistent with APPsi/e>>APPsi/E4>APPsi/E3 (**Fig. 2a-j**). In the cortex and hippocampus, Aβ burden in APPsi/e mice was higher than APPsi/E3 mice (p<0.01 in cortex and p<0.0001 in hippocampus) and APPsi/E4 mice (p=0.0538 in cortex and p<0.01 in hippocampus) (**Fig. 2d, i**). Further analysis revealed that the burden in APPsi/E4 mice trended higher than APPsi/E3 mice (p=0.0591 in cortex and p<0.05 in hippocampus) (**Fig. 2e, j**). Analysis of Aβ levels by sequential buffer-extraction and ELISA confirmed that both formic acid (FA)-soluble and SDS-soluble amyloid burden was highest in the parental APPsi/e mice, with Aβ42 species dominating over Aβ40 (**Fig. 2k-n**). Surprisingly, in the FA-soluble fraction, we observed that both APPsi/e and APPsi/E4 had higher biochemical levels of Aβ42 and Aβ40 relative to APPsi/E3 (**Fig. 2k-l**; p<0.0001 in FA fraction, p<0.05 in SDS fraction), in spite of APPsi/E4 showing limited parenchymal deposits. FA-associated Aβ42 levels were slightly higher in APPsi/e mice compared to APPsi/E4 (**Fig. 2k**; p<0.05). In the SDS-soluble fraction, APPsi/e mice had higher Aβ loads compared to APPsi/E3 and APPsi/E4 mice (**Fig. 2m-n**; p<0.0001 relative to APPsi/E3 and p<0.01 relative to APPsi/E4 for Aβ42; p<0.01 relative to APPsi/E3 and p<0.05 relative to APPsi/E4 for Aβ40). Overall, these results indicate that relative to mouse apoE, human apoE3 and apoE4 suppressed the deposition of diffuse Aβ with apoE3 showing somewhat greater suppression.

**Figure 2.**
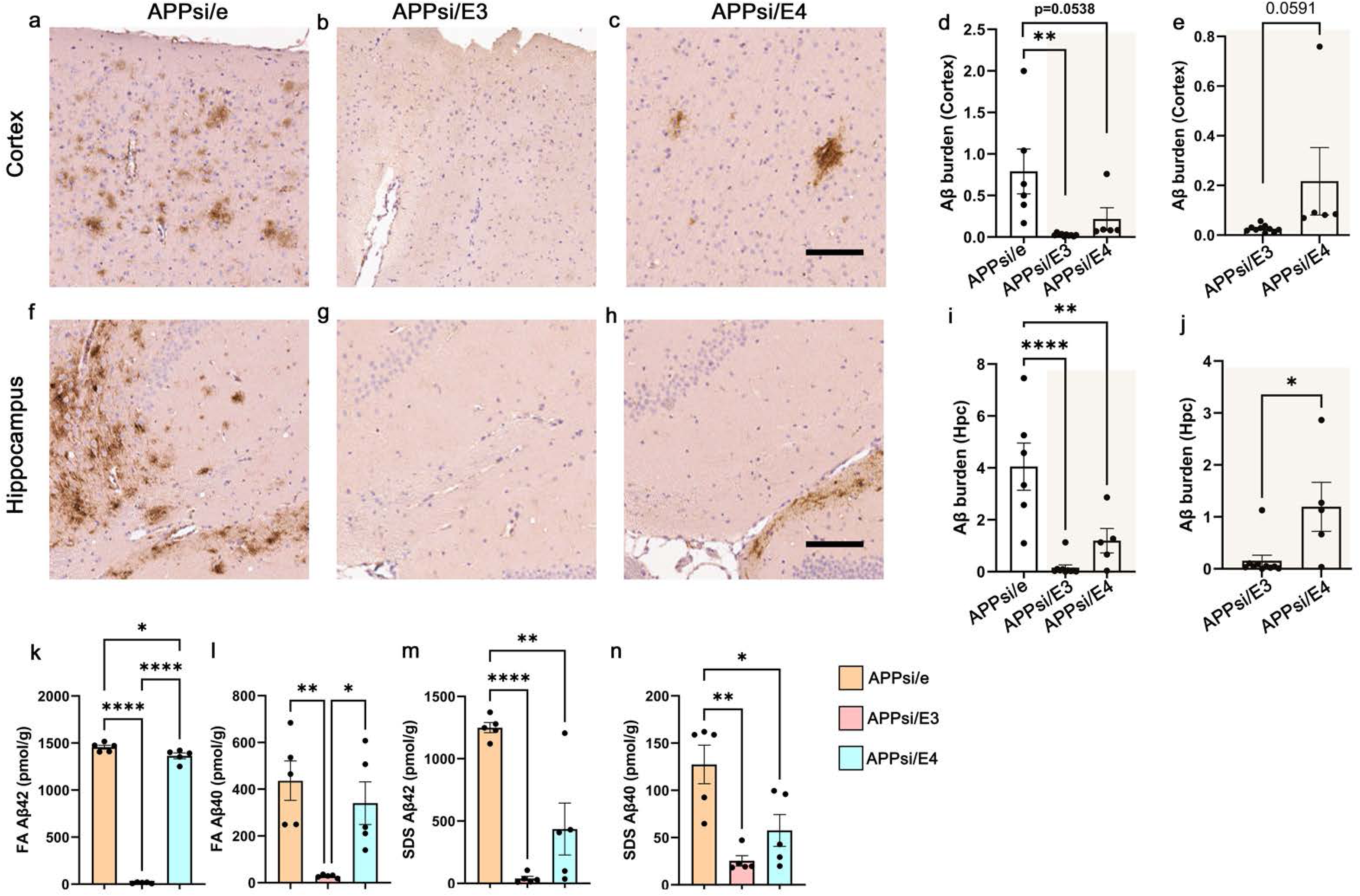
Aβ burden in aged APPsi mice expressing mouse and human apoE isoforms. **a-j.** Representative FFPE sections showing Aβ plaques in the cortex (a-c) and hippocampus (f-h) of 21-mo old bigenic APPsi mice that are homozygous for mouse *Apoe*, or human *APOE3* and *APOE4*. Three sections from each mouse were used for quantitation of Aβ burden in the cortex (d, e) and hippocampus (i, j). Analysis was done among all three strains (d, i) or within the human *APOE* genotypes (e, j). **k-n** Serially extracted brain lysates were used to evaluate biochemical levels of Aβ40 and Aβ42 in the formic acid (FA)-associated fraction (k, l) and SDS-associated fraction (m, n) using Aβ end-specific sandwich ELISA. n=6 mice/group. 1-way Anova with Tukey’s test (d, I, k, l, m, n) and unpaired 2-tailed t test (e, j). ****p<0.0001, **p<0.01, *p<0.05. Scale bar, 100 µm.

### Cerebral seeding mitigates the differential effects of human *APOE3* and *APOE4*

Next, we examined whether apoE isoforms can modify the seeded induction of Aβ aggregation. Intracerebral amyloid seeding leading to Aβ plaque deposition in the brain shares many properties with the prion templating process, including substantial acceleration in the onset and progression of Aβ aggregation (reviewed in [37]). In a previous study, we had shown that neonatal intracerebral seeding of APPsi mice with homogenates prepared from the brains of AD patients accelerated the onset of amyloid deposition in these mice producing robust diffuse Aβ deposits by 12-mo of age [24]. Following passaging and amplification of the human brain-derived seeds in APPsi mice, brains of seeded APPsi were collected at 13-mo of age to generate the inoculum for seeding a secondary cohort of neonatal APPsi mice. We injected these mouse-passaged amyloid seeds into the cerebral ventricles of APPsi/E4 and APPsi/E3 mice and aged the mice to 12-mo to test whether apoE isoforms had differential effect of seeded propagation of Aβ. A cohort of similarly-seeded 12-mo old APOEtr mice were used as controls. At 12-mo of age, the seeded APPsi/E4 and APPsi/E3 mice showed similar levels of cortical and hippocampal Aβ burden (**Fig. 3a-d**) that were much higher than that of the unseeded 21-mo old APPsi/e mice (compare **Fig. 3a-d to Fig. 2d-j**). None of the Aβ-homogenate seeded 12-mo old APOEtr mice developed any Aβ deposits (**Fig. 3a-d**). ThioS staining also confirmed that the morphology of the plaques in all the strains of seeded mice was of the diffuse type (**Additional File 1:Fig. S2c**), consistent with our earlier report that seeded APPsi mice develop mostly diffuse Aβ deposits [24]. One notable aspect in this case is that neither seeding-mediated acceleration nor the presence of apoE4 resulted in emergence of dense-core plaques. Thioflavin S staining revealed that vascular deposition was limited in these seeded 12-mo old bigenic APPsi x APOEtr mice (**Additional File 1:Fig. S2c**). Analysis of Aβ levels by sequential buffer-extraction and ELISA confirmed that SDS-soluble and FA-soluble amyloid biochemical load was similar between the APPsi/E4 and APPsi/E3 mouse lines examined and the predominant form was Aβ42 (**Fig. 3e-h**). These biochemical levels were consistent with our observation in brain homogenate seeded APPsi/e mice reported earlier [24]. Thus, the solubility characteristics of Aβ40 and Aβ42 were not altered by introduction of human *APOE* alleles. We also examined if different apoE isoforms led to unique patterns of astrocytosis and microgliosis (**Additional File 1:Fig. S3**). Quantification of the burden of these markers revealed elevated GFAP levels in seeded APPsi/E4 mice relative to APPsi/E3 (p<0.05 in cortex and p<0.01 in hippocampus) and APOE4tr mice (p<0.05 in cortex and hippocampus) (**Additional File 1:Fig. S3a-d**). We did not observe any differential induction of microgliosis in these mice (**Additional File 1:Fig. S3e-h**). Overall, these observations indicate that any differential effects of human *APOE* alleles on Aβ were mitigated by accelerating Aβ deposition through a templating process.

**Figure 3.**
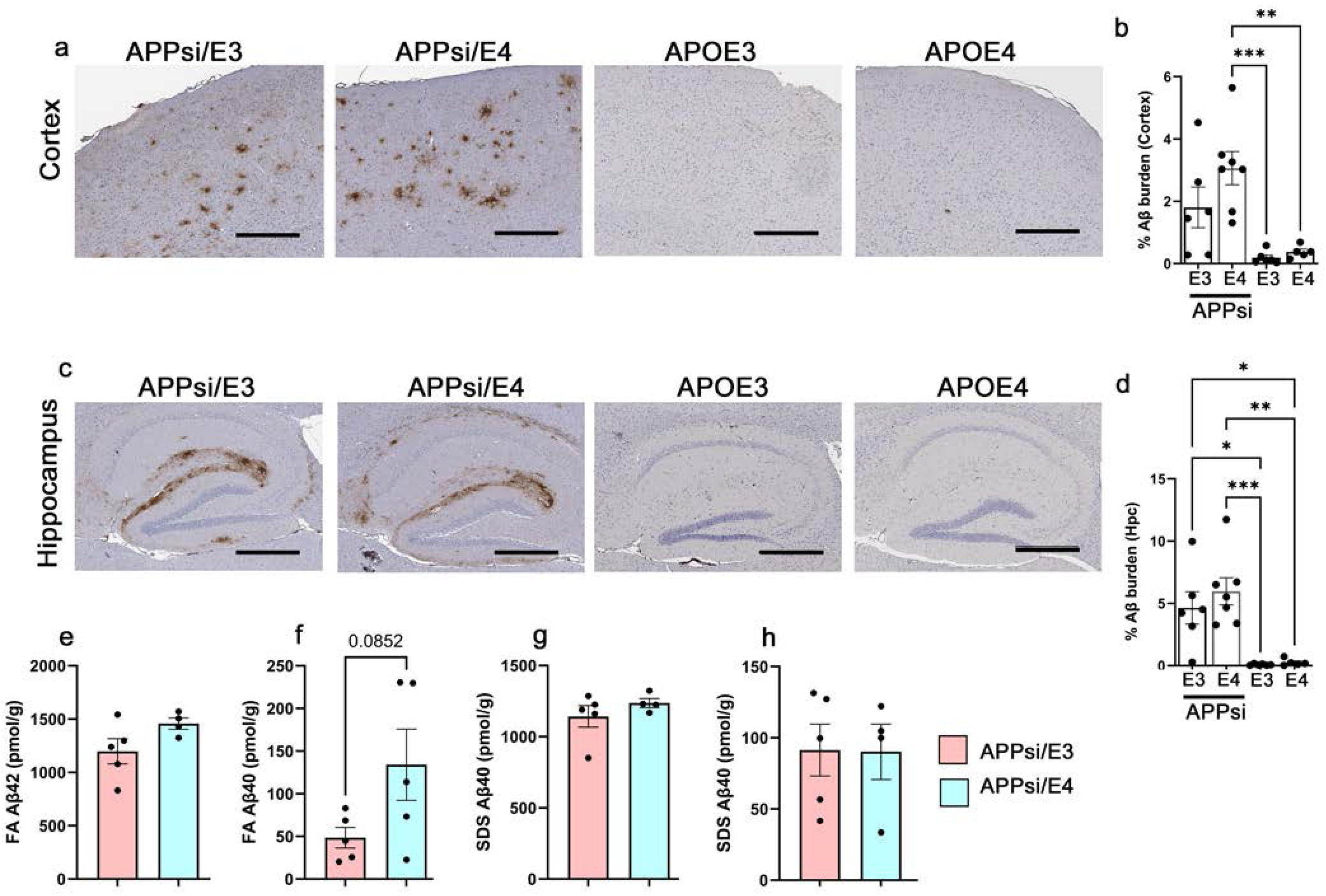
Aβ burden in intracerebrally-seeded APPsi mice expressing mouse and human apoE isoforms. **a-d.** Representative FFPE sections showing Aβ plaques in the cortex (a) and hippocampus (c) of 12-mo old APPsi/*APOE3*, APPsi/*APOE4*, APOE3tr and APOE4tr mice that were seeded in the cerebral ventricles as neonates. Three sections from each mouse were used for quantitation of Aβ burden in the cortex (b) and hippocampus (d). **e-h.** Serially extracted brain lysates were used to evaluate biochemical levels of Aβ40 and Aβ42 in the FA-associated fraction (e, f) and SDS-associated fraction (g, h). n=6 for human *APOE* mice. 1-way Anova with Tukey’s test (b, d) and unpaired t-test (e-h). **p<0.01, *p<0.05. Scale bar, 500 µm.

We wanted to determine if there was evidence of differential interaction of apoE protein with these diffuse Aβ deposits in seeded APPsi/E3 and APPsi/E4 mice. First, we stained three serial sections with antibodies to Aβ and two apoE antibodies, one recognizing both mouse and human proteins and the other specific for human apoE4 (**Fig. 4**). We found that both human apoE3 and apoE4 were copiously associated with Aβ deposits (arrows, **Fig. 4a**). Co-immunofluorescence analysis further confirmed this observation (**Fig. 4b**).

**Figure 4.**
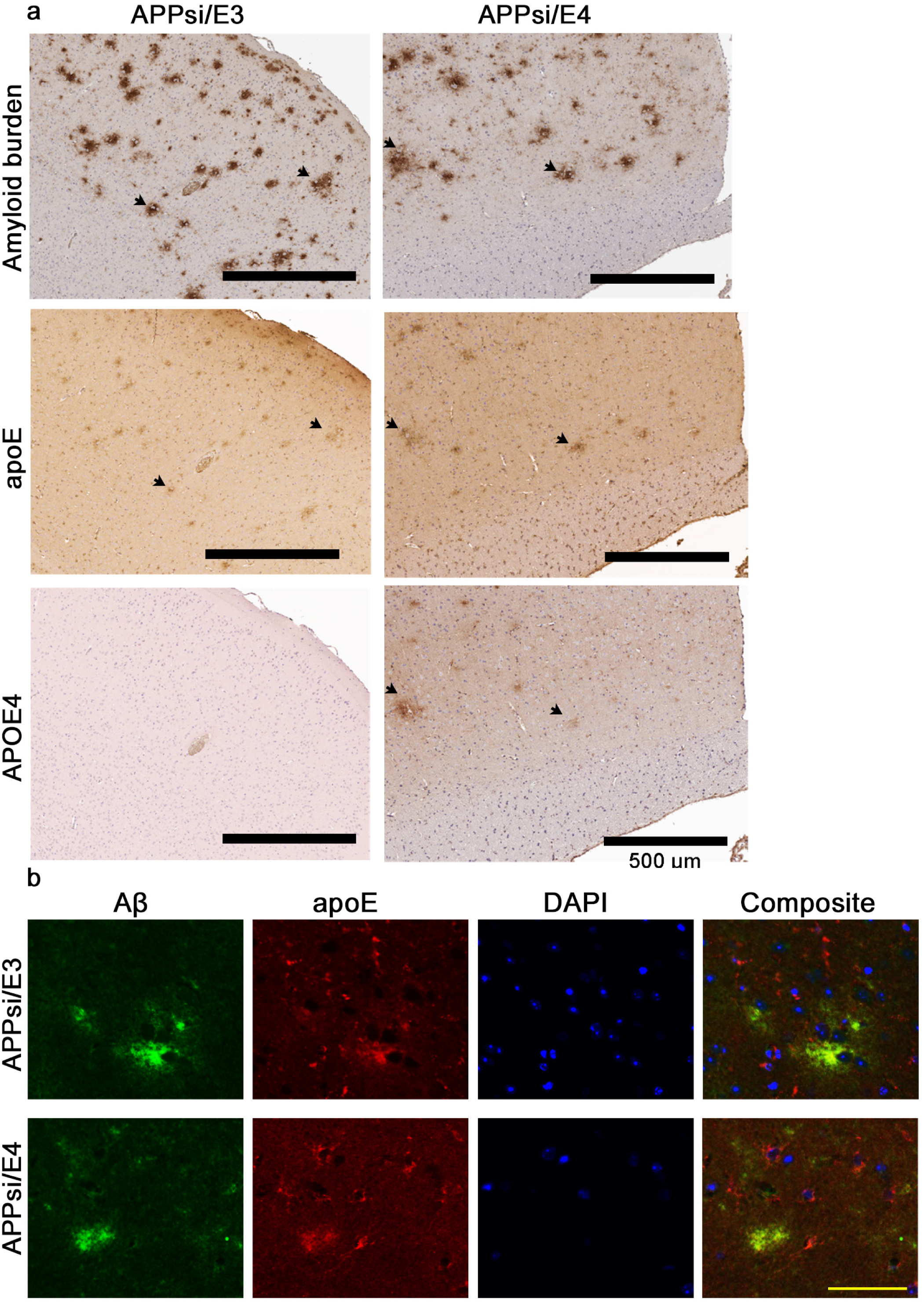
apoE-Aβ colocation in intracerebrally-seeded APPsi mice expressing mouse and human apoE isoforms. **a.** Representative FFPE sections of bigenic APPsi mice that are homozygous for human *APOE3* (12-mo) and *APOE4* (12-mo). Three serial sections were stained with pan Aβ antibody, pan apoE antibody and an apoE4-specific antibody. Arrows denote apoE staining on putative Aβ plaques. Scale bar, 500 µm. **b.** Co-immunofluorescence showing apoE (Alexa fluor 594nm) decorated Aβ deposits (Alexa fluor 488nm). DAPI is used as nuclear counterstain. Scale bar, 50 µm.

### Human apoE isoforms suppress Aβ deposition relative to mouse apoE in a model of dense-cored and fibrous Aβ

The recently developed SAA-APP knock-in mouse model develops fibrillar Aβ deposits within the cortex and hippocampus, with homozygous mice developing deposits by 5 months of age and the amyloid burden increasing progressively with advancing age [26]. Although a recent study examined the influence of *APOE* genotype on amyloid deposition in a similar APP knock-in model [38], a study of the effects of *APOE* allelic variation on amyloid deposition in the SAA-APP model has not been reported to date. To characterize this new knock-in model, we crossed SAA-APP mice to humanized APOEtr mice to generate double homozygous animals for each of four possible apoE isoforms; mouse *Apoe* or human *APOE2*, *APOE3*, or *APOE4*. Mice aged 5-mo or 8-mo of age were analyzed for neuropathology. We confirmed that the full-length APP and C terminal fragment levels in these different strains were equivalent (**Additional File 1:Fig. S4a-d**). Quantification of multiple sections from each animal of each genotype revealed a rank order of amyloid deposition burden in SAA-APP mice with mouse apoE being the highest followed by SAA-APP/E4 mice with SAA-APP/E2 and SAA-APP/E3 mice showing the least amyloid deposition (**Fig. 5a-h**; p<0.0001 for mouse apoE vs all human apoE variants in cortex and hippocampus). Thus, similar to our findings in APPsi mice, the presence of human apoE variants, particularly apoE3, strongly suppresses the deposition of cored-fibrillar Aβ deposits.

**Figure 5.**
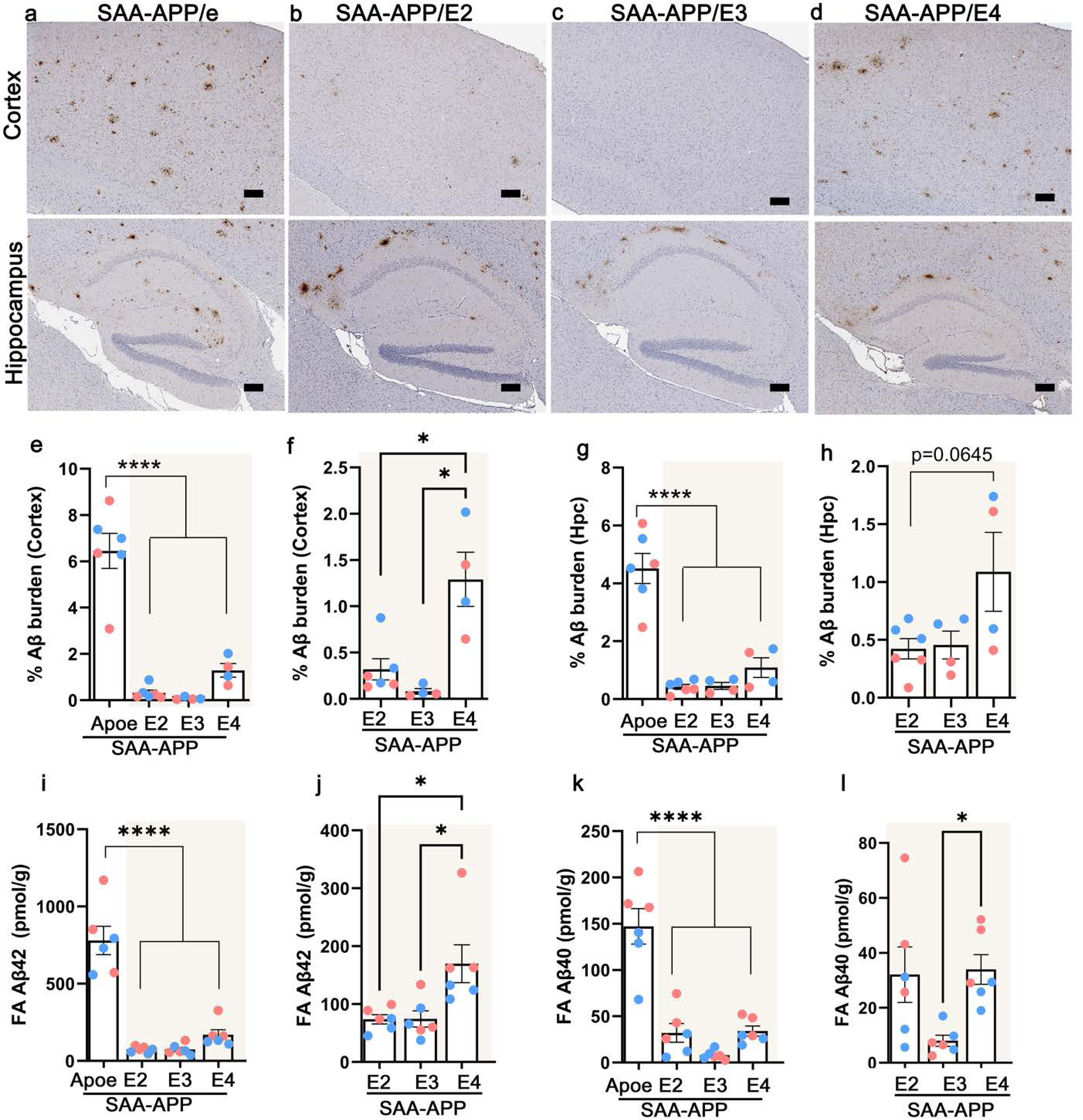
Aβ burden in SAA-APP mice expressing mouse and human apoE isoforms. **a-d.** Representative FFPE sections showing Aβ plaques in the cortex and hippocampus of 8-mo old bigenic SAA-APP mice that are homozygous for mouse *Apoe*, or human *APOE2*, human *APOE3* and *APOE4*. **e-h.** Three sections from each mouse were used for quantitation of Aβ burden in the cortex (e, f) and hippocampus (g, h). Analysis was done among all four strains (e, g) or within the human *APOE* genotypes (f, h). n=6 mice/group, blue indicates male mice and pink indicates female mice. Scale bar, 500 µm. 1-way Anova with Tukey’s test. **i-l.** FA-extracted brain lysates were used to evaluate biochemical levels of insoluble Aβ42 (i-j) and Aβ40 (k-l). Analysis was done among all four strains (I, k) or within the human *APOE* genotypes (j, l). n=6 mice/group. 1-way Anova with Tukey’s test. ****p<0.0001, ***p<0.001, **p<0.01, *p<0.05.

Analysis within the human *APOE* sub-group revealed that SAA-APP/E4 mice had higher cortical deposits relative to SAA-APP/E3 and SAA-APP/E2 mice at 8-mo of age (**Fig. 5f**; p<0.05). In the hippocampus, SAA-APP/E4 mice showed a trend towards higher burden compared to SAA-APP/E2 and SAA-APP/E3 mice (**Fig. 5h**; p=0.0645). We confirmed presence of dense-cored plaques in these mice using Thioflavin S staining (**Additional File 1:Suppl. Fig. S5**). ELISA analysis of biochemical levels of soluble and insoluble Aβ revealed SAA-APP/e mice had substantially higher levels of both Aβ40 and 42 relative to SAA-APP mice bearing human *APOE* alleles (**Fig. 5i-k**; p<0.0001). In general, FA-associated Aβ40 levels were lower than the corresponding Aβ42 levels, with the highest levels observed in SAA-APP/e mice, relative to all the human APOE variants (**Fig. 5k**; p<0.0001). Within the human *APOE* genotypes, *APOE4* mice had higher insoluble Aβ42 than *APOE3* or *APOE2* (**Fig. 5j**; p<0.05) and the FA-associated Aβ40 levels in SAA-APP/E4 were higher than SAA-APP/E3 (p<0.05) but not in SAA-APP/E2 mice (**Fig. 5l**). These findings generally confirm the pathological assessments, as the levels of insoluble Aβ40 and Aβ42 were highest in the brains of SAA-APP/e relative to the others. In addition, our data indicates that there is a preference for the Aβ42 species in this model, irrespective of the apoE isoform present.

We observed similar outcomes in a second cohort of mice aged 5-7 months (**Fig. S6a**). The cortical Aβ burden of 5-mo SAA-APP/e mice was significantly higher than 5.4-mo old SAA-APP/E4 (range 5.3-5.7-mo), 6.4-mo old SAA-APP/E3 (range 5.5-7.3-mo) and 6.8-mo old SAA-APP/E2 (6.3-7.3-mo) mice (**Additional File 1:Fig. S6a-b;** p<0.0001 relative to all human genotypes). Within-group analysis showed that cortical Aβ burden in *APOE4* mice was higher than *APOE3* (p<0.01) and *APOE2* (p<0.05) mice (**Additional File 1:Fig. S6c**). In the hippocampus, all four strains examined had equivalent Aβ burden, with higher trends observed in SAA-APP/e mice (**Additional File 1:Fig. S6e-f**). This finding was also consistent with increased neurodegenerative disease-associated microglial transcriptome signatures in the SAA-APP/e mice relative to mice bearing the human *APOE* alleles, while the levels were comparable within the three human *APOE* allele containing SAA-APP mice (**Additional File 1:Fig. S6g-i**). Collectively, these findings indicate that the presence of human *APOE* allelic variants in mouse models of amyloidosis strongly suppresses amyloid deposition relative to mouse *Apoe*.

Next, we examined the relative levels of apoE protein in the brains of these mice, finding no significant differences in the soluble apoE protein between the different human apoE derivatized SAA-APP strains (**Additional File 1:Fig. S4e-g**). We found lower levels of mouse apoE, which could be due to the detection threshold of the antibody and the type of apoE particles (**Additional File 1:Fig. S4g**). We then confirmed the presence of apoE binding to individual plaques across the strains. All strains of SAA-APP mice, whether bearing mouse apoE or human apoE, demonstrated Aβ-apoE co-staining, though there were subtle variations that could be related to the overall Aβ burden (**Fig. 6**).

**Figure 6.**
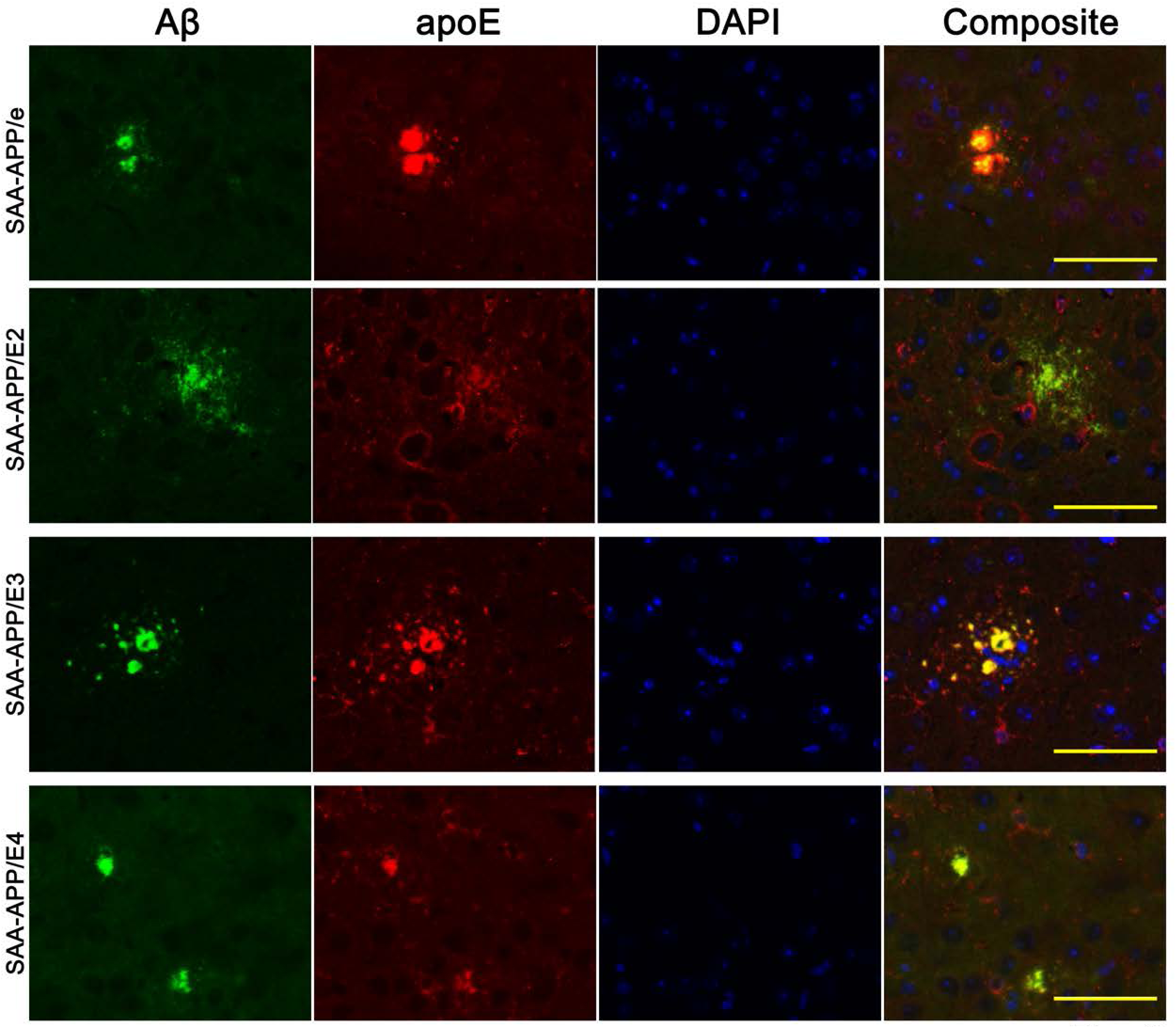
apoE-Aβ colocation in SAA-APP expressing mouse and human apoE isoforms. Co-immunofluorescence showing apoE (Alexa fluor 594nm) decorated Aβ deposits (Alexa fluor 488nm) in 8-mo old SAA-APP mice carrying mouse *Apoe* or human *APOE* alleles. DAPI is used as nuclear counterstain. Scale bar, 50 µm.

To determine the influence of *APOE* allelic variants on inflammatory changes that occur in SAA-APP mice as amyloid is deposited, we stained brain sections from the 8-mo old cohorts with antibodies to GFAP and Iba1 (**Additional File 1:Fig. S7**). The cortical GFAP was largely proportional to amyloid burden in the SAA-APP mice, i.e., the degree of immunostaining followed the pattern *Apoe*>>A*POE4*∼*APOE3*∼*APOE2* (**Additional File 1:Fig. S7a-c**), while there were no significant alterations in the hippocampus (**Additional File 1:Fig. S7d-e**). Iba-1 labeled microgliosis showed more complicated patterns, with *APOE4* showing higher Iba-1 levels than *APOE2*- and *Apoe*-bearing SAA-APP mice in the cortex and hippocampus (**Additional File 1:Fig. S7f-j**). For these experiments, we included controls in the form of age-matched APP-WT knock-in mice crossed to APOE3tr and APOE4tr mice (Ct, **Additional File 1:Fig. S7b, d, g, i**). Using silver staining and immunohistochemistry, we first confirmed that, as expected, 8-mo old APP-WT/e, APP-WT/E3 and APP-WT/E4 mice lacked amyloid deposits (**Additional File 1:Fig. S8a-b**). The brains of these APP-WT mice (with human *APOE3* or *APOE4* alleles) also showed low levels of GFAP and Iba1 immunoreactivity (**Additional File 1:Fig. S8c-d**) and thus were collapsed as a single group to determine baseline gliosis in Fig. S7. Compared to these baseline APP-WT cohort, the brains of SAA-APP/e mice had higher levels of cortical GFAP reactivity (p<0.01) and Iba1 (p<0.05) (**Additional File 1:Fig. S7b, g**). Both SAA-APP/E3 and SAA-APP/E4 mice had higher Iba-1 relative to WT-APP mice with *APOE3* and *APOE4* (**Fig. S7g, i**), though there were no similar changes in GFAP (**Fig. S7b, d**). Within the human alleles, we seldom saw any statistically significant changes, except in microgliosis in SAA-APP/E3 mice relative to SAA-APP/E2 mice (**Additional File 1:Fig. S7j**; p<0.05) and suggestive trends between SAA-APP/E4 mice vs. SAA-APP/E2 mice (**Additional File 1:Fig. S7h, j**; p=0.0529 in cortex and p=0.0620 in hippocampus). Overall, it appeared that astrogliosis strongly correlated with amyloid deposition, and microgliosis correlated with presence of *APOE4* and *APOE3* alleles.

## Discussion

*APOE* genotype is the major genetic risk factor of AD and one of its key properties is regulation of the onset and progression of Aβ deposition [39]. Here, using three distinct paradigms, we show that (1) relative to mouse *Apoe*, presence of human *APOE* alleles suppress the deposition of both diffuse and cored-fibrillar Aβ deposits in transgenic mouse models, with *APOE3* having a greater impact compared to *APOE4*, (2) the suppressive activities of human *APOE* alleles were mitigated by inducing diffuse Aβ deposition through seeding where both *APOE3* and *APOE4* show comparable amyloidogenicity, (3) *APOE4* was slightly more amyloidogenic than *APOE3* in a naturally progressing model of diffuse amyloidosis, and (4) the presence of human *APOE* alleles does not impact the innate morphology of deposits that arise naturally or through seeding.

The APPsi model, through natural progression and when seeded intracerebrally primarily develops diffuse Aβ deposits, with seeding accelerating the appearance of diffuse amyloid significantly [24,25]. In comparing the amyloid burden in the brains of the unseeded APPsi/E3 and APPsi/E4 mice we found that they showed similar burden that was much much lower than the amyloid burden of age-matched APPsi/e mice. In cortex, the burden of amyloid trended higher in unseeded APPsi/E4 mice but did not achieve statistical significance. Notably, APPsi/E4 mice also had a higher burden of vascular amyloid compared to APPsi/E3 mice, consistent with a previous report showing that presence of apoE4 shifts the Aβ from parenchymal to the vascular compartment [27]. Measurements of Aβ levels in unseeded mice revealed consistently higher level of Aβ42 and Aβ40 in APPsi/E4 mice, indicating that the presence of human apoE4 was less suppressive than that for human apoE3. It remains to be established whether high levels of CAA found in APPsi/E4 mice contributed to the higher levels of FA-associated Aβ compared to the corresponding levels in the APPsi/E3 mice.

The amyloid burden in seeded APPsi x APOEtr mice was much higher than that of unseeded mice and similar to what we previously reported for APPsi mice (bearing mouse apoE) seeded with human AD brain homogenates [24], suggesting that inducing amyloid deposition by seeding overcame the suppressive effects of human apoE on Aβ deposition. This finding may shed some additional insight into the potential mechanism by which the presence of human apoE suppresses Aβ deposition. Using different models, two independent studies showed that apoE was critical in the early assembly of Aβ deposits rather than affecting plaque growth, with apoE4 showing a stronger effect on neuritic plaque development [34,35]. We suggest that seeding negates the early effects of apoE on Aβ aggregation to induce deposition even in the presence of a strong suppressor such as human apoE3.

The question of why mouse apoE is a weaker suppressor of amyloid deposition (both diffuse and dense-cored) relative to human apoE remains unknown. It is possible that mouse apoE is less able to clear human Aβ peptides or early Aβ oligomers than human apoE through direct or indirect mechanisms [40–43]. Mouse apoE may bind to Aβ to prime aggregation more effectively [4], or have a different preference for lipid or lipid transporter molecules that influence Aβ clearance from interstitial fluids [17]. Mouse and human apoE may have differential interactions with canonical apoE interactors such as heparan sulphate [44,45], which could act as a substrate for the initial deposition of Aβ (reviewed in [46]) or have differential capacity to stimulate microglial uptake and clearance of Aβ [42]. Lastly, there could be differences in the post-translational modification of mouse and human apoE that affect their interaction with Aβ [47]. While it could be a mix of these different mechanisms, the several fold differences in amyloidogenicity between mouse and human apoE (∼60 times higher than human apoE3 and 5x times higher than human apoE4 in the SAA-APP/e mice) could be harnessed to provide molecular insights into the structural moieties on apoE that directly influence Aβ deposition. There is a future prospect of translating these insights into therapeutic outcomes by guiding the discovery of agents that antagonize apoE-receptor or apoE-ligand interactions to achieve lower amyloid burden in susceptible individuals.

Recent publications have argued that non-neuronal apoE, once secreted, can bind to Aβ and facilitate its compaction [48]. Despite our clear observation that human apoE3 and apoE4 were associated with diffuse Aβ deposits, we observed no obvious remodeling of diffuse deposits into cored deposits in the time frame of our experiment. In the 21-mo old naturally progressing APPsi cohort and the intracerebrally-seeded 12-mo old APPsi cohorts, presence of the human apoE isoforms did not influence the innately diffuse Aβ deposition phenotype nor did it influence the location of the deposits. In the SAA-APP cohort that has dense-cored and fibrous deposits, we observed Aβ deposits more consistently in the cortex. While the cortical Aβ plaques were evenly distributed throughout all layers of the cortex, the hippocampal deposits were assembled at the boundary with the corpus callosum. Morphologically, the deposits were similar between the four different SAA-APP cohorts, irrespective of the apoE isoform present (**Additional File 1:Fig. S9**). Across all genotypes, there were large course-grained neuritic deposits, smaller dense deposits, and numerous small punctate deposits, most often observed in SAA-APP/e mice and SAA-APP/E4 mice (**Additional File 1:Fig. S9**). We observed significant levels of apoE protein decorating the plaques in all of the SAA-APP and seeded APPsi strains. Thus, while we observed an association of apoE with both diffuse and cored deposits, we found no evidence that this association influenced the morphological appearance or anatomical location of Aβ pathology.

In AD patients, there is a wide diversity of parenchymal Aβ deposit morphology. An early study in post-mortem AD brains reported that the relative proportion of diffuse, fibrillar and dense-cored plaques was 31, 49 and 20% and the proportion of neuritic pathology in these three types of deposits were 24, 82 and 76% [49]. Data on resilience factors in AD have shown that not all individuals with Aβ deposits will develop dementia associated with AD. Indeed, specific phenotypic characteristics, such as presence of dense-cored plaques and neuritic pathology are associated with pathological progression into dementia [13]. Relative to the diffuse plaques, the dense cored plaques are associated with neuritic malformations and reactive gliosis (reviewed in [50]), factors that could enhance neurodegenerative outcomes. Older human patients also accumulate a small proportion of ‘burnt-out’ plaques which are composed of exclusively of dense cores with no neuritic components which are thought to represent the end-stage of Aβ plaque life cycle [51]. Based on the progressive morphological states of plaques found in human patients – diffuse plaques appearing earlier than dense-cored and burnt-out plaques - it is speculated that plaques originate as diffuse deposits that are then compacted during plaque growth phase into dense-core plaques and evolving finally to burnt-out morphology [52–54]. However, other studies have strongly suggested that amyloid plaque types arise de novo, perhaps mediated via a seeding process, without any progressive maturation involved [25,55–57]. Our observations in the naturally progressing APPsi/E4 mice would suggest that even a strong pro-amyloidogenic factor such as apoE4, that has been shown in earlier studies to be a key molecule in early phases of plaque seeding [34], is not sufficient to alter the innately diffuse deposits to dense-cored morphology. Based on this finding, we suggest that plaques arise de novo based on ambient genetic and metabolic conditions, and that plaque life cycle may not necessarily go through maturation phases.

### Limitations

Our study does not provide data on behavioral assessment of these mice. In addition, future insights into how different apoE-lipid particles determine Aβ strain selection and formation of specific types of deposits can provide information on the differences observed in these models.

### Conclusions

Our study uses three different new paradigms to establish how different apoE isoforms influence the progression of Aβ plaque pathology in naturally aging animals and in intracerebrally-seeded animals.

## Abbreviations

Aβ: amyloid β
AD: Alzheimer’s disease
*APOE*: Human Apolipoprotein E allele
APOEtr: mice homozygous for human *APOE2*, *APOE3* or *APOE4* alleles via targeted replacement of the murine *Apoe* gene
*Apoe*: Murine Apolipoprotein E allele
apoE: Apolipoprotein E protein
APPsi/e: PrP.APPsi mice homozygous for murine *Apoe*
APPsi/E3, APPsi/E3H: PrP.APPsi mice homozygous for human *APOE3*
APPsi/E4, APPsi/E4H: PrP.APPsi mice homozygous for human *APOE4*
APPsi/E3h: PrP.APPsi mice heterozygous for human *APOE3*
APPsi/E4h: PrP.APPsi mice heterozygous for human *APOE4*
FFPE: formalin-fixed paraffin-embedded
GFAP: Glial fibrillary acidic protein
Iba-1: Ionized calcium binding adaptor molecule 1
Ntg: Nontransgenic for APP/Aβ
SAA-APP/e: hAbetaSAA mice with murine *Apoe*
SAA-APP/E2: hAbetaSAA mice homozygous for human *APOE2*
SAA-APP/E3: hAbetaSAA mice homozygous for human *APOE3*
SAA-APP/E4: hAbetaSAA mice homozygous for human *APOE4*
WT-APP/e: B6J hAbeta mice with murine *Apoe*
WT-APP/E3: B6J hAbeta mice homozygous for human *APOE3*
WT-APP/E4: B6J hAbeta mice homozygous for human *APOE4*

## Author contributions

PC, DRB: Conceptualization, Data curation, Formal analysis, Funding acquisition, Investigation, Methodology, Project administration, Software, Supervision, Validation, Visualization, Writing - original draft, Writing - review & editing GX, PS: Data curation, Formal analysis, Investigation, Methodology, Validation, Visualization, Writing - review & editing PB, CG, SF, QV, KNM, QL: Investigation; Formal Analysis

## Ethics approval

Animals use in this study was approved by the Institutional Animal Care and Use Committee at the University of Florida.

## Competing Interests

The authors declare no competing interests exist.

## Additional Information: Supplementary Material

Supplementary Figures S1 – S9; Supplementary Table S1, S2, S3.

## Availability of data and materials

The datasets supporting the conclusions of this article are included within the article and its additional files.

## Funding Declaration

The work was supported by NIA RF1AG057933 (DRB, PC) and NIA R01AG055798 (PC).

## Acknowledgements

We thank Zhi Huang for help with *APOE* genotyping and Selma S Brkic and Amanda Lopez for neuropathological analysis.

## Notes

### Competing Interest Statement

The authors have declared no competing interest.

