## Supplementary Figures and Tables for "Human *APOE* allelic variants suppress the formation of diffuse and fibrillar Aβ deposits relative to mouse *Apoe* in transgenic mouse models of Alzheimer amyloidosis"

**Additional File 1: Supplementary Material**

**Supplementary Table 1. Antibodies used in the study.**

| Antibody | Ligand | Host | Catalog # | Vendor | Dilution | Assay |
|---|---|---|---|---|---|---|
| 33.1.1 | Aβ - N terminus | Mouse | N/A | Todd Golde | 1:13,000 | IHC, IF |
| GFAP | GFAP | Rabbit | Z0334 | Dako | 1:1000 | IHC |
| Iba-1 | Iba-1 | Rabbit | 019-19741 | Wako | 1:1000 | IHC |
| apoE | apoE | Goat | AB947 | EMD Millipore | 1:1000 | IHC, IF |
| APOE4 | Human apoE4 | Mouse | MABN43 | EMD Millipore | 1:1000 | IHC |
| UBi-1 | Ubiquitin | Mouse | MCA-Ubi-1 | EnCor | 1:1000 | IHC |
| CT20 | APP C terminus | Rabbit | N/A | Todd Golde | 1:1000 | Western |
| apoE | apoE | Goat | AB947 | EMD Millipore | 1:1000 | Western |
| Actin | β-actin | Mouse | A 5441 | Sigma | 1:1000 | Western |
| 2.1.3 | Aβ42 | Mouse | N/A | Todd Golde | 25μg/ml | ELISA |
| 13.1.1 | Aβ40 | Mouse | N/A | Todd Golde | 25μg/ml | ELISA |
| 33.1.1-HRP | Aβ - N terminus | Mouse | N/A | Todd Golde | 1:1000 | ELISA |
| Donkey anti-goat IgG, Alexa Fluor 594 | Mouse IgG | Donkey | A-11058 | Life Technologies | 1:500 | IF |
| Donkey anti-Mouse IgG, Alexa Fluor 488 | Mouse IgG | Donkey | A-21202 | Life Technologies | 1:500 | IF |

**Supplementary Table 2. Neuropathological burden analysis of parenchymal Aβ deposits and CAA score in silver-stained sections from 21-mo old APPsi and homozygous APPsi/*APOE* mice.** Details of mice used for calculating amyloid score and CAA score in Figure 1. The following criteria was used for Amyloid scoring: 3 = >50; 2 = 10-50; 1 = <10; 0 = 0-1 discrete deposit(s) per section. The following criteria was used for CAA scoring: 3 = numerous stained parenchymal vessels per section; 2 = frequent stained vessels per section; 1 = limited number of stained vessels per section; 0 = no stained vessels per section.

| Genotype | Animal ID | Sex | Age | Amyloid Score | CAA score |
|---|---|---|---|---|---|
| APPsi B6n4 | 1062923-1 | M | 21.0 | 2 | 3 |
| APPsi B6n4 | 1066532-1 | M | 21.0 | 3 | 3 |
| APPsi B6n5 | 1101469-1 | F | 21.0 | 2 | 2 |
| APPsi B6n5 | 1101469-2 | F | 21.0 | 3 | 3 |
| APPsi B6n5 | 1113538-2 | F | 21.0 | 2 | 3 |
| APPsi B6n5 | 1113539-1 | M | 21.0 | 3 | 2 |
| APPsi B6n5 | 1113540-1 | F | 21.0 | 3 | 2 |
| APPsi B6n5 | 1126228-2 | M | 21.0 | 2 | 3 |
| APPsi B6n5 | 1126235-1 | F | 21.0 | 3 | 3 |
| APPsi B6n5 | 1162467-1 | M | 21.0 | 2 | 3 |
| APPsi B6n5 | 1162470-1 | F | 21.0 | 2 | 3 |
| APPsi B6n5 | 1162470-2 | F | 21.0 | 3 | 3 |
| APPsi B6n6 | 1210993-1 | M | 21.0 | 3 | 2 |
| APPsi/APOE3+/+ | 1198249-1 | M | 20.0 | 0 | 1 |
| APPsi/APOE3+/+ | 1044023-1 | F | 21.0 | 0 | 1 |
| APPsi/APOE3+/+ | 1053163-1 | F | 21.0 | 0 | 0 |
| APPsi/APOE3+/+ | 1198232-1 | F | 21.0 | 0 | 0 |
| APPsi/APOE3+/+ | 1198232-2 | F | 21.0 | 0 | 1 |
| APPsi/APOE3+/+ | 1198232-3 | F | 21.0 | 0 | 2 |
| APPsi/APOE3+/+ | 1201977-4 | F | 21.0 | 0 | 1 |
| APPsi/APOE3+/+ | 1204248-4 | F | 21.0 | 1 | 1 |
| APPsi/APOE3+/+ | 1204248-5 | F | 21.0 | 0 | 1 |
| APPsi/APOE3+/+ | 1198235-1 | M | 21.0 | 0 | 2 |
| APPsi/APOE3+/+ | 1201919-2 | M | 21.0 | 0 | 0 |
| APPsi/APOE4+/+ | 1024757-3 | F | 19.4 | 0 | 3 |
| APPsi/APOE4+/+ | 1024763-1 | M | 19.4 | 1 | 2 |
| APPsi/APOE4+/+ | 1011810-4 | F | 20.8 | 0 | 2 |
| APPsi/APOE4+/+ | 1167757-2 | M | 20.8 | 1 | 3 |
| APPsi/APOE4+/+ | 1011811-2 | M | 20.9 | 1 | 3 |
| APPsi/APOE4+/+ | 1011811-3 | M | 20.9 | 1 | 3 |
| APPsi/APOE4+/+ | 1172114-4 | M | 20.9 | 1 | 1 |
| APPsi/APOE4+/+ | 1184293-2 | F | 21.0 | 1 | 3 |

**Supplementary Table 3. Neuropathological burden analysis of parenchymal Aβ deposits and CAA score in silver-stained sections from 21-mo old heterozygous APPsi/APOE mice.** Details of mice used for calculating amyloid score and CAA score in Figure 1 and Figure S1. The following criteria was used for Amyloid scoring: 3 = >50; 2 = 10-50; 1 = <10; 0 = 0-1 discrete deposit(s) per section. The following criteria was used for CAA scoring: 3 = numerous stained parenchymal vessels per section; 2 = frequent stained vessels per section; 1 = limited number of stained vessels per section; 0 = no stained vessels per section.

| Genotype | Animal ID | Sex | Age | Amyloid Score | CAA score |
|---|---|---|---|---|---|
| APPsi/APOE3+/- | 1162464-1 | F | 21.0 | 0 | 1 |
| APPsi/APOE3+/- | 1169180-1 | F | 21.0 | 1 | 2 |
| APPsi/APOE3+/- | 1201912-1 | F | 21.0 | 1 | 2 |
| APPsi/APOE3+/- | 1201977-1 | F | 21.0 | 2 | 1 |
| APPsi/APOE3+/- | 1201977-2 | F | 21.0 | 2 | 1 |
| APPsi/APOE3+/- | 1204248-1 | F | 21.0 | 2 | 2 |
| APPsi/APOE3+/- | 1204248-2 | F | 21.0 | 2 | 2 |
| APPsi/APOE3+/- | 1204248-3 | F | 21.0 | 3 | 2 |
| APPsi/APOE3+/- | 1162465-1 | M | 21.0 | 1 | 1 |
| APPsi/APOE3+/- | 1201915-1 | M | 21.0 | 3 | 2 |
| APPsi/APOE3+/- | 1201915-4 | M | 21.0 | 2 | 2 |
| APPsi/APOE3+/- | 1201919-1 | M | 21.0 | 1 | 1 |
| APPsi/APOE3+/- | 1201919-3 | M | 21.0 | 1 | 1 |
| APPsi/APOE3+/- | 1204249-2 | M | 21.0 | 2 | 2 |
| APPsi/APOE4+/- | 1121213-1 | F | 20.9 | 0 | 3 |
| APPsi/APOE4+/- | 1121213-2 | F | 20.9 | 2 | 1 |
| APPsi/APOE4+/- | 1160920-3 | F | 20.9 | 2 | 1 |
| APPsi/APOE4+/- | 1160920-4 | F | 20.9 | 2 | 2 |
| APPsi/APOE4+/- | 1114748-1 | M | 20.9 | 2 | 1 |
| APPsi/APOE4+/- | 1121215-1 | M | 20.9 | 2 | 3 |
| APPsi/APOE4+/- | 1184293-4 | F | 21.0 | 3 | 2 |

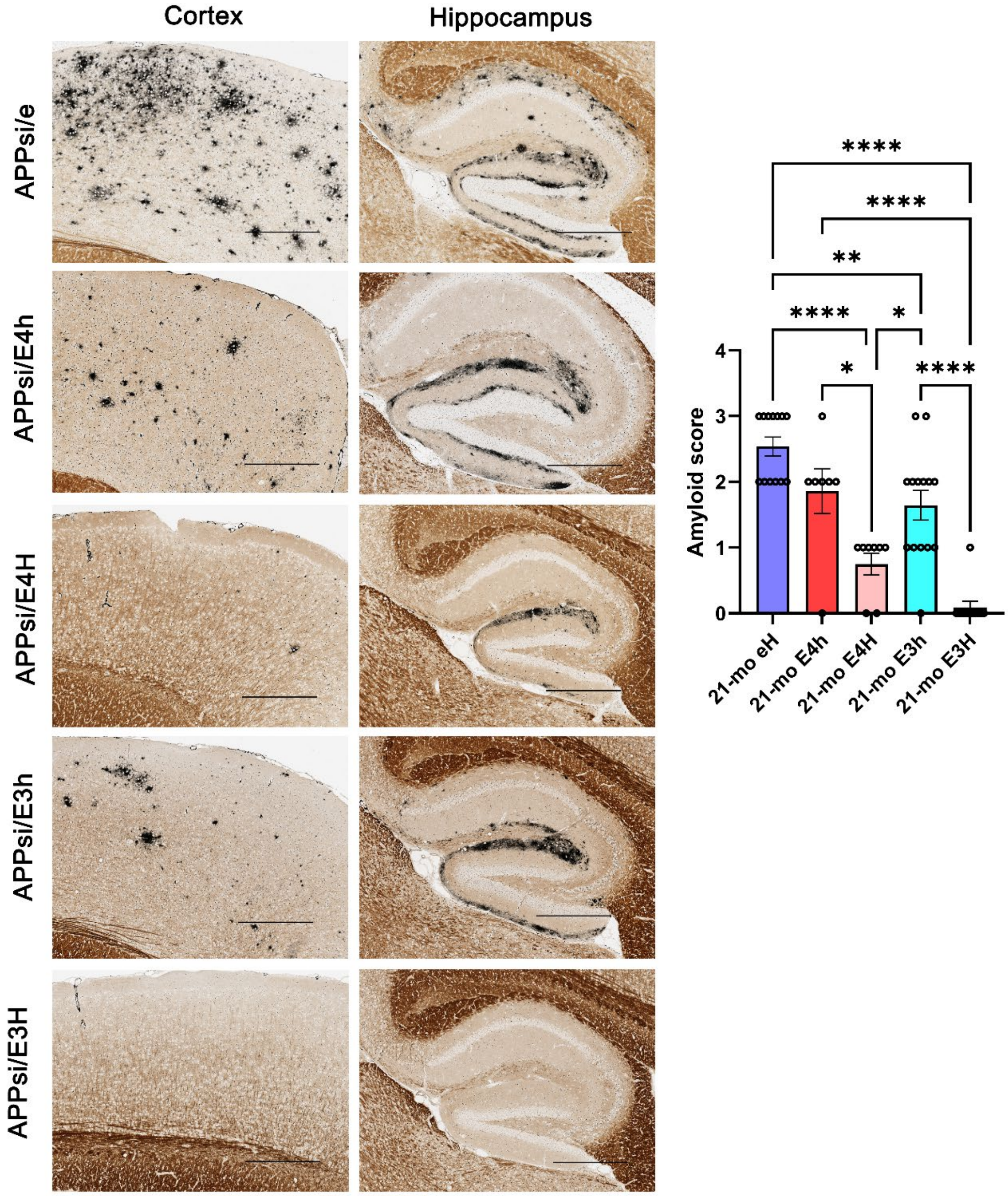

Suppl. Fig. S1

**Suppl. Fig. S1. Mouse apoE is dominant over human apoE in 21-mo old APPsi strain.**

Representative silver-stained sections from APPsi homozygous for mouse *Apoe* (APPsi/e), human *APOE3* (APPsi/E3H) and human *APOE4* (APPsi/E4H) as well as APPsi mice heterozygous for human *APOE3* (APPsi/E3h) or human *APOE4* (APPsi/E4h). Amyloid score was calculated using neuropathological burden criteria and plotted as a graph. 1-way Anova, ****p<0.0001, **p<0.01, *p<0.05. n= 13 for APPsi/e, n = 11 for APPsi/E3H, n=8 for APPsi/E4H mice, n = 14 for APPsi/E3h, n = 7 for APPsi/E4h. Scale bar, 100 µm.

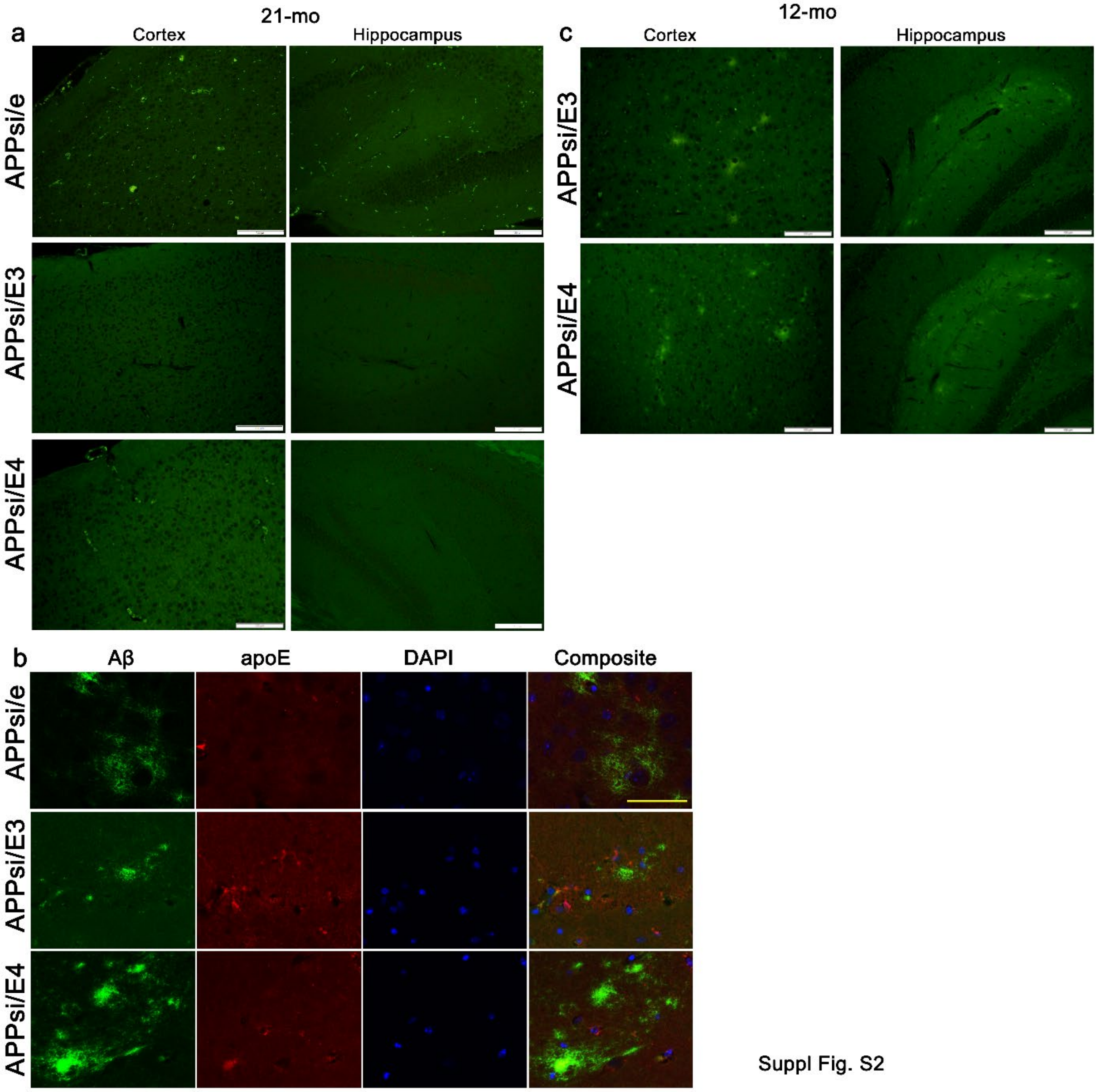

**Suppl. Fig. 2. Evaluation of cored plaques in 21-mo old APPsi mice and intracerebrally-seeded 12-mo old APPsi mice.**

**a** Representative Thioflavin S-stained images from the cortex and hippocampus of 21-mo old bigenic APPsi mice that are homozygous for mouse *Apoe*, or human *APOE3* and *APOE4*. **b** Co-immunofluorescence of FFPE sections on 21-mo old mouse brain sections was done using Alexa fluor 468nm for a pan Aβ antibody and Alexa fluor 594nm for a pan apoE antibody. DAPI is used as a nuclear stain. **c** Representative Thioflavin S-stained images from the cortex and hippocampus of 12-mo old bigenic APPsi mice that are homozygous for human *APOE3* and *APOE4* following intracerebral seeding as neonates. Scale bar, 100 μm. n=3-6 mice/group.

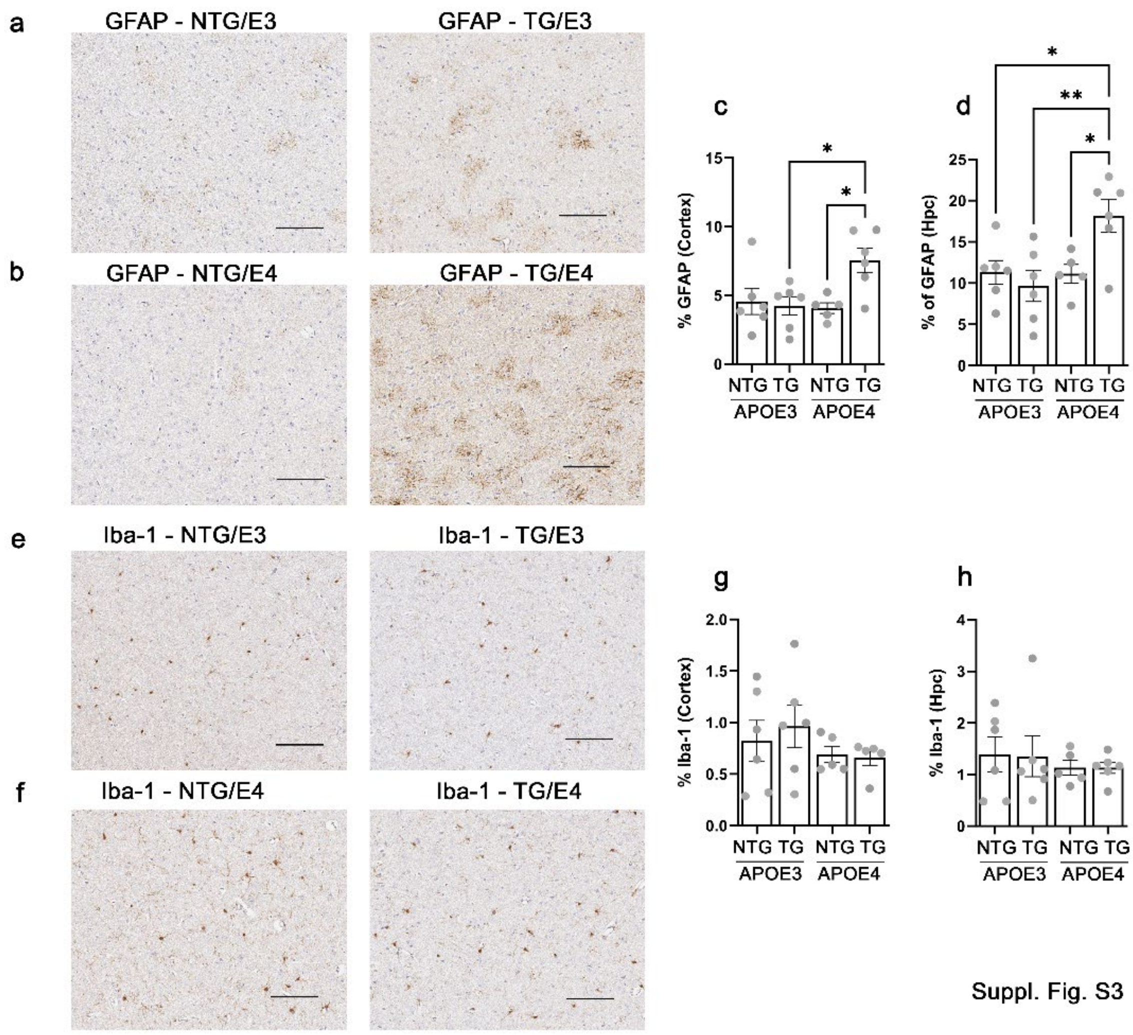

**Suppl. Fig. S3. Gliosis patterns in 12-mo old intracerebrally-seeded APPsi mice expressing mouse and human apoE isoforms**.

**a,b** Representative FFPE sections showing GFAP-labeled astrocytes in the cortex of 12-mo old APPsi mice carrying human *APOE3* or human *APOE4* alleles (TG) or APOE3tr and APOE4 tr mice (NTG) seeded intracerebrally as neonates. **c,d** Quantification of staining shown in cortex and hippocampus. **e,f** Representative FFPE sections showing Iba-1-labeled microglia in the cortex of 12-mo old APPsi mice carrying human *APOE3* or human *APOE4* alleles (TG) or *APOE3tr* and *APOE4tr* mice (NTG) seeded intracerebrally as neonates. **g,h** Quantification of staining shown in cortex and hippocampus. Scale bar, 100 µm. 1-way Anova. **p<0.01, *p<0.05. n=6 mice/group.

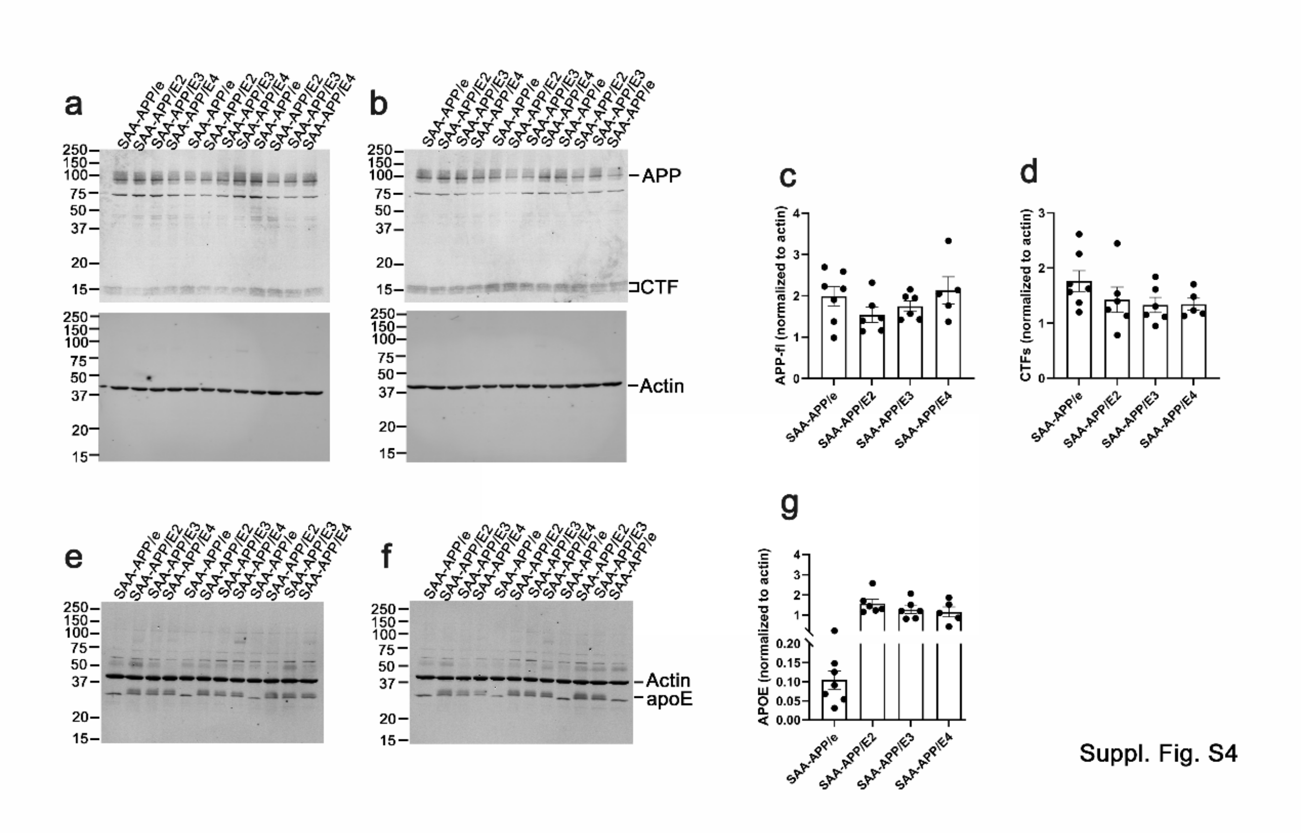

**Suppl. Fig. S4. Evaluation of APP and apoE levels in SAA-APP mice**.

**a-g** Immunoblots showing full length APP (APPfl) and C-terminal fragments (CTF) (a-d) and apoE protein levels (e-g) in 5-7-mo old bigenic SAA-APP mice that are homozygous for mouse *Apoe*, or human *APOE2*, human *APOE3* and *APOE4*. Numbers on the left indicate molecular weight markers in kDa. Quantitation of indicated protein bands, normalized to house-keeping gene actin, shown in right hand panels (c, d, g). n=5-7 mice/group.

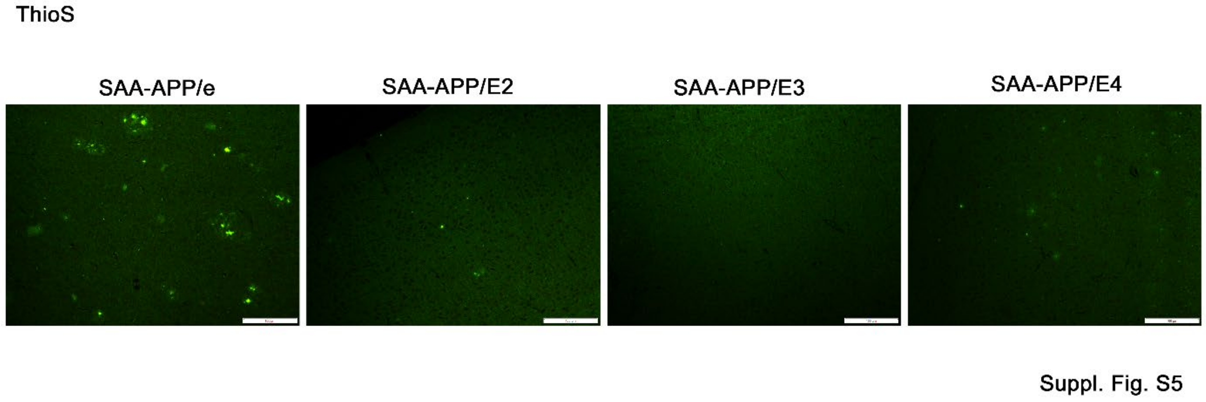

**Suppl. Fig. S5. Evaluation of cored plaques in 8-mo old SAA-APP mice**.

Representative Thioflavin S-stained images from the cortex of 8-mo old bigenic SAA-APP mice that are homozygous for mouse *Apoe*, or human *APOE2*, human *APOE3* and *APOE4*. Scale bar, 100 µm. n=6 mice/group.

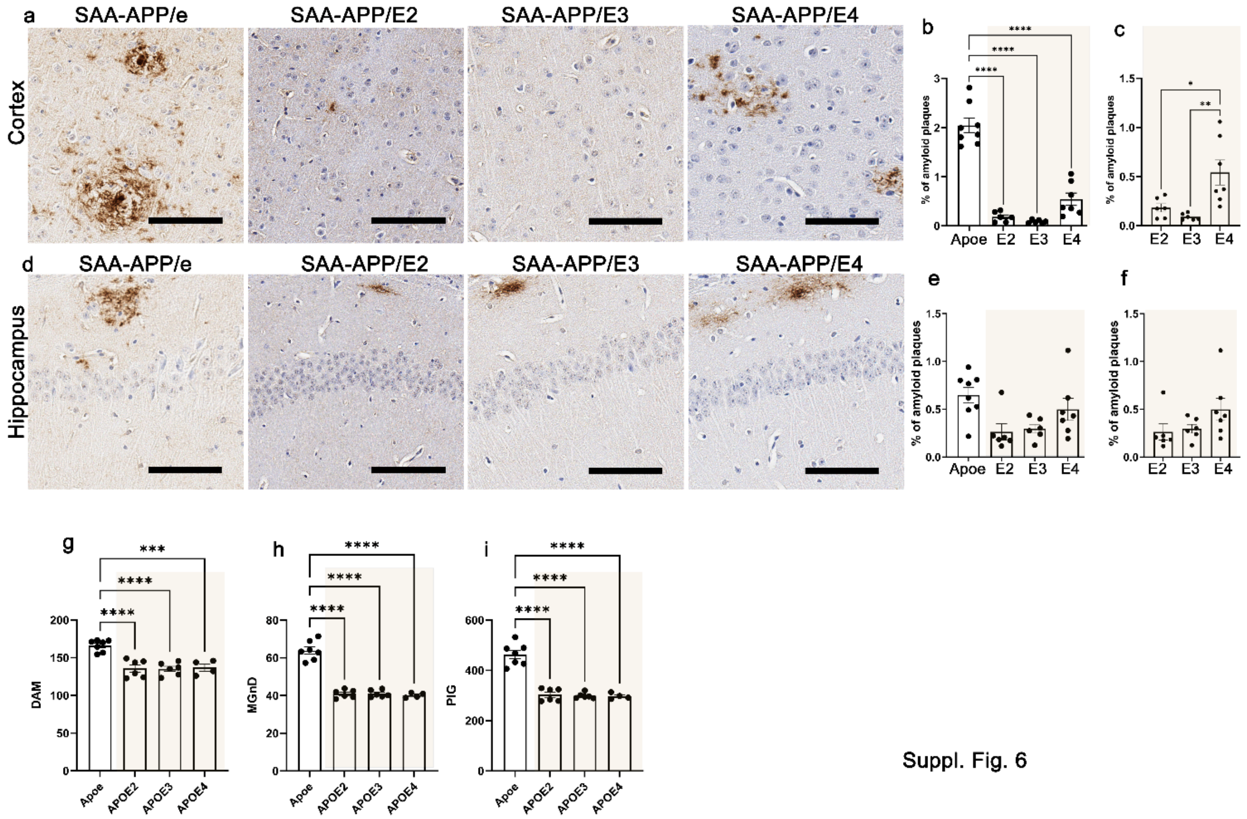

Suppl. Fig. 6

**Suppl. Fig. S6. Aβ burden in young SAA-APP mice expressing mouse and human apoE isoforms**.

**a,d** Representative FFPE sections showing Aβ plaques in the cortex and hippocampus of young bigenic SAA-APP mice that are homozygous for mouse *Apoe* 5-mo), or human *APOE2* (~6.8-mo), human *APOE3* (~6.4-mo) and *APOE4* (~5.5-mo). **b,c; e,f** 3 sections from each mouse were used for quantitation of Aβ burden in the cortex (b, c) and hippocampus (e, f). Analysis was done among all four strains (b, e) or within the human *APOE* genotypes (c, f). Scale bar, 100 μm. 1-way Anova. n=5-6 mice/group. **g-i** AD-typical transcriptomic signatures signifying neurodegenerative processes have been imputed from NanoString RNA analysis. Analysis shows the prevalence of damage-associated microglia (DAM, g), neurodegenerative microglia (MGnD, h) and plaque-induced gees (PIG, i) in these mice. n=4-7 mice/group.

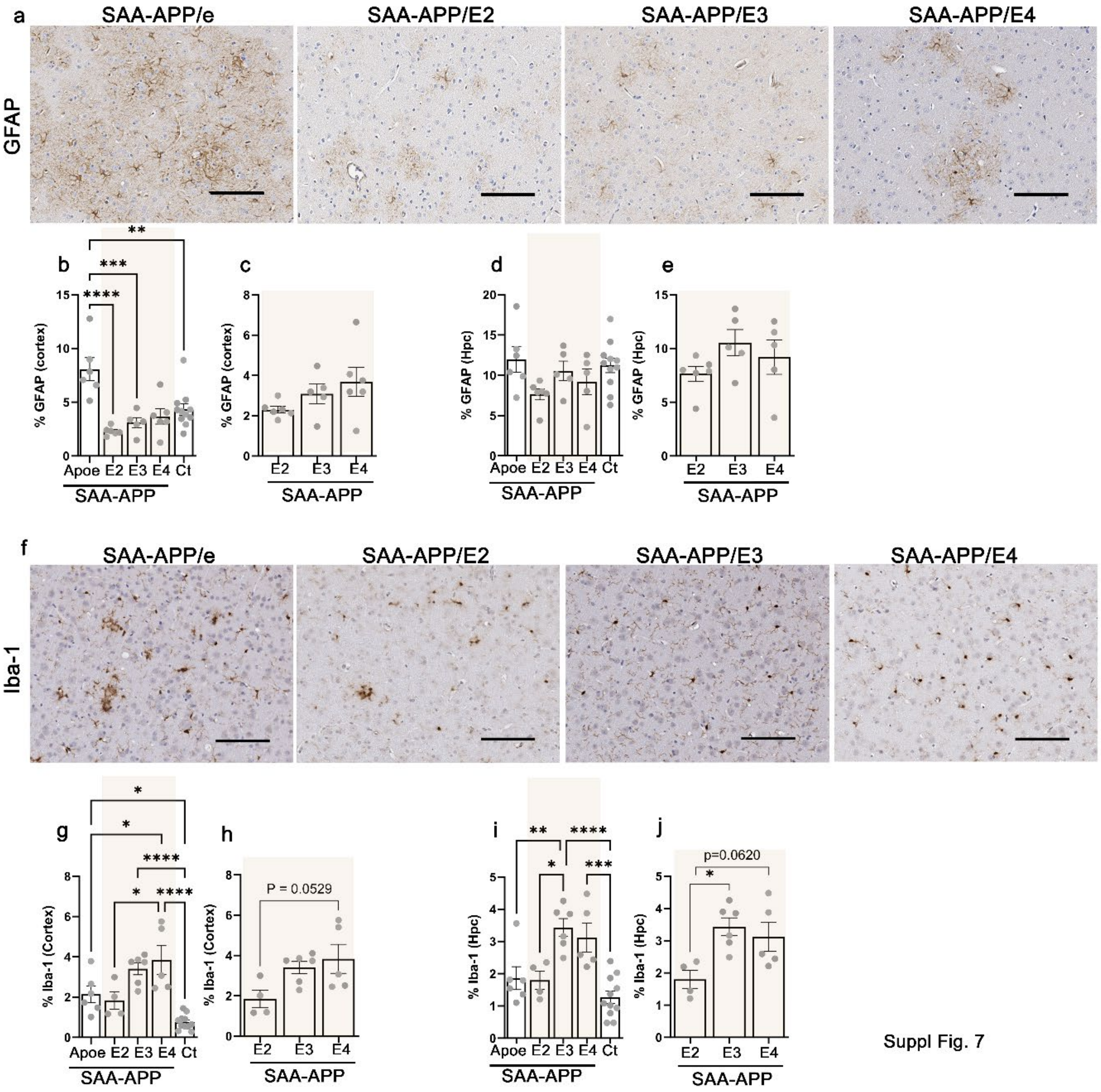

**Suppl. Fig. S7. Gliosis patterns in SAA-APP mice expressing mouse and human apoE isoforms**.

**a** Representative FFPE sections showing GFAP-labeled astrocytes in the cortex of 8-mo old SAA-APP mice carrying mouse *Apoe*, human *APOE2*, *APOE3* or human *APOE4* alleles. **b-e** Quantification of staining shown in cortex and hippocampus across all strains (b, d) or within human *APOE* cohorts (c, e). Nontransgenic (NTG) control (Ct) denote WT-APP mice that are homozygous for *APOE3* and *APOE4*. **f** Representative FFPE sections showing Iba-1-labeled microglia in the cortex of 8-mo old SAA-APP mice carrying mouse *Apoe*, human *APOE2*, *APOE3* or human *APOE4* alleles. **g-j** Quantification of staining shown in cortex and hippocampus across all strains (g, i) or within human *APOE* cohorts (h, j). Nontransgenic (NTG) control (Ct) denote WT-APP mice that are homozygous for *APOE3* and *APOE4*. Scale bar, 100 µm. ****p<0.0001, ***p<0.001, **p<0.01, *p<0.05. 1-way Anova. n=6 mice/group.

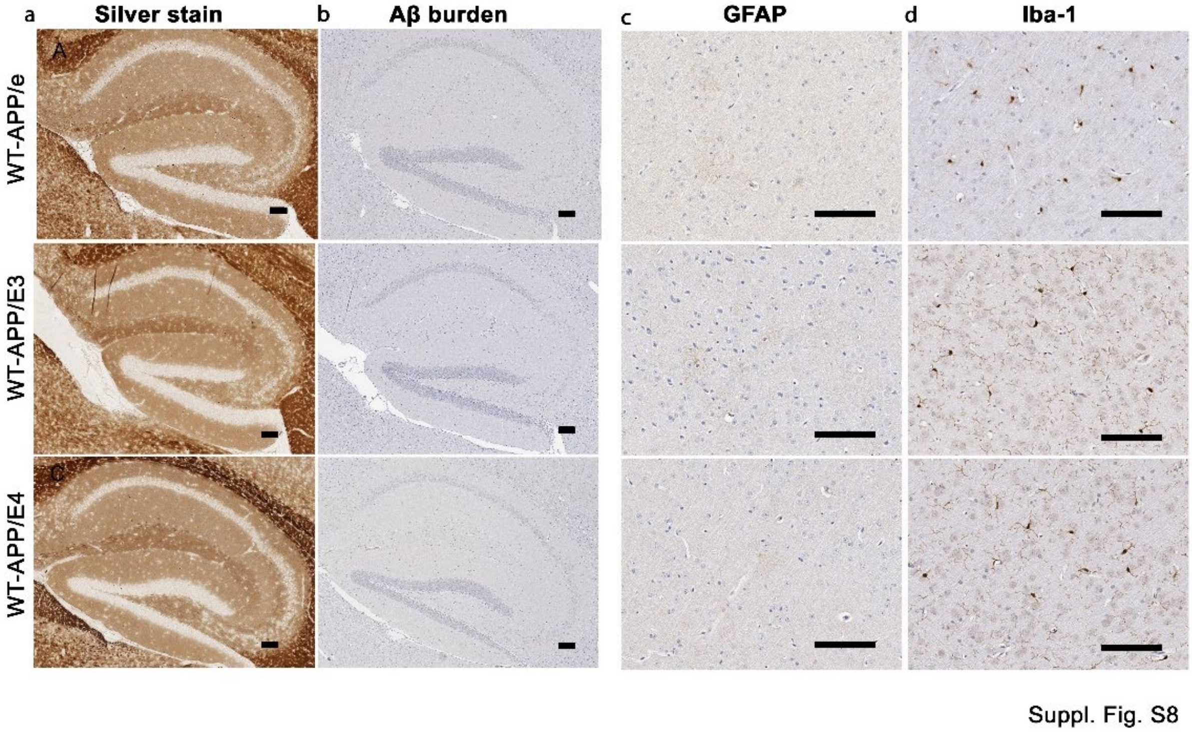

**Suppl. Fig. S8. Characterization of aged WT-APP mice expressing mouse and human apoE isoforms**.

**a,b** Representative stained images from the brains of 12-14-mo old WT-APP mice that carry mouse *Apoe* or human *APOE3* and *APOE4*. Aβ has been visualized using Campbell-Switzzer silver stain (a) and a pan Aβ antibody (b). **c,d** Astrocytosis (GFAP, c) and microgliosis (Iba-1, d) shown from these same mice. n=4 mice/group. Scale bar, 100 μm.

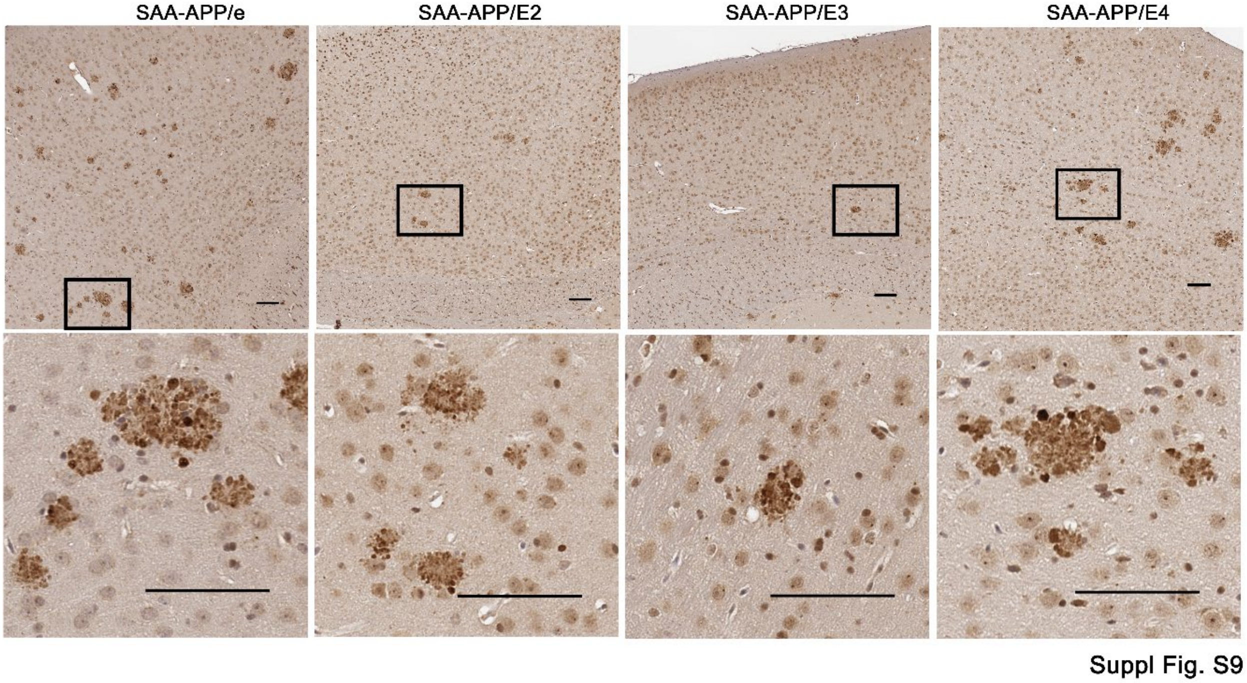

**Suppl. Fig. 9. Summary of plaque morphology in SAA-APP strains**.

Representative images from Ubiquitin antibody stained cortex from cortical 8-mo SAA-APP mice carrying different apoE isoforms. Individual boxed areas on top panel have been magnified to show morphological characteristics of amyloid deposits in each cohort (bottom panel). Scale bar, 100 µm. n=6 mice/group.
